# A B1 SINE element within the mouse *Malat1* long noncoding RNA acts as an internal ribosome entry site mediating micropeptide translation

**DOI:** 10.64898/2026.02.11.705401

**Authors:** Wen Xiao, Tanja Hann, Zion Perry, Vijaya Pandey, Jeffrey Cheng, James Wohlschlegel, Anna Marie Pyle, Douglas L. Black

## Abstract

In most cells, *Malat1* long noncoding RNA localizes to the nucleus where it affects splicing and chromatin function. In mouse neurons, *Malat1* is exported to the cytoplasm where it is translated to generate the M1 micropeptide. Here we characterize an internal ribosome entry site (IRES) required for M1 translation that is sufficient to induce cap-independent expression of a downstream ORF. *In vivo* chemical probing and structural modeling defined a 3-stem-loop RNA secondary structure that encompasses the active IRES segment and coincides with a mouse B1 SINE repetitive element. In affinity purification experiments, a minimal IRES/B1 RNA selectively bound ribosomal subunits, translation factors, and the known IRES cofactors Rack1 and hnRNP A2/B1. Depletion of Rack1 and hnRNP A2/B1 inhibited IRES-dependent translation of the reporter, and expression of the endogenous M1 peptide. Our study identifies an unexpected functional unit hidden within a widely studied long noncoding RNA.

**Teaser:** A repetitive sequence element embedded in a long noncoding RNA drives hidden micropeptide production in mammalian cells.

## Introduction

Many cellular RNAs originally classified as noncoding have been found to contain very short open reading frames that encode micropeptides (*1–5*). We recently showed that the long noncoding RNA *Malat1* in mouse encodes at least one micropeptide (*6*). *Malat1* is broadly expressed as an abundant nuclear RNA that is studied for its effects on splicing and chromatin (*7–11*). We found that mouse *Malat1* accumulates in the cytoplasm of developing cortical neurons where it is packaged with proteins into granules and transported to neuronal processes. This cytoplasmic *Malat1* is bound by ribosomes and translated to produce a peptide called M1. Neuronal M1 translation is stimulated by depolarization, and depletion of *Malat1* or blocking M1 translation leads to enhanced expression of synaptic proteins such as synaptophysin, indicating that M1 may act to dampen physiological responses(*6*). *Malat1* RNA also affects synaptogenesis in the developing mouse cortex, and is implicated in fear extinction memory(*12*, *13*). Thus, in addition to its canonical roles in the nucleus, *Malat1* in mouse neurons is also an mRNA affecting synaptic function.

*Malat1* has features that distinguish it from typical mRNAs. It is an RNA Pol II transcript with a standard 7-methyl guanosine cap(*14*), but is predominantly unspliced and is not polyadenylated(*15–17*). The 3’ end of *Malat1* RNA is created by an unusual processing reaction. The 3’ terminus of the primary transcript contains a tRNA-like structure, the Masc RNA, that is released from the transcript through cleavage by RNAse P(*15*). The resulting 3’ end of *Malat1* folds into a stable triple helical structure that protects it from exonucleolytic decay, and was shown to support translation of reporter mRNAs that lack polyA tails(*16–18*). Multiple nucleotides within *Malat1* have been found to carry modifications including 6-methyl adenosine (m6A), pseudouridine, and 5-methyl cytosine(*19–21*). Although their stoichiometry is not known, m6A nucleotides are found in the synaptically localized RNA, raising the possibility that they could modulate the expression of its encoded peptide.

Internal ribosome entry sites (IRES) are specialized RNA sequences that induce cap-independent translation initiation at sites within the body of an RNA. These discrete functional units are found in many viruses and some cellular mRNAs, and generally act to recruit ribosomal subunits to a downstream AUG initiation codon without the action of eIF4E protein binding at the mRNA cap. IRES’s are diverse in structure and are classified by their varying cofactor dependencies.

We noted that the initiation codon for the M1 peptide in mouse is 739 nucleotides downstream of the Malat1 Cap and is preceded by numerous other initiation codons. This raised the question of whether the M1 peptide is translated via some noncanonical initiation mechanism. We show here that the M1 ORF is preceded by a structured RNA segment that acts as an internal ribosome entry site to stimulate its translation. This sequence coincides with a B1 short interspersed element (SINE) and is sufficient to recruit ribosomal subunits to an RNA. These studies describe how an RNA molecule widely studied as a nuclear noncoding transcript can also be translated through a noncanonical mechanism.

## Results

### Translation of the mouse *Malat1* M1 ORF requires an upstream regulatory sequence

To identify *Malat1* sequences upstream of the M1 ORF that affect its translation, we made a reporter gene with GFP fused in frame to the C-terminus of M1 (Fig. 1A). This contained the entire *Malat1* coding sequence including the sequences required for *Malat1* 3’ processing and was driven by the CMV promoter. The plasmid was transfected into mouse N2A neuroblastoma cells and translation of the M1 ORF was assayed by GFP expression (Fig. 1A). As seen previously(*6*), GFP expression was dependent on the M1 translation initiation codon at *Malat1* nucleotide 740 (Fig. 1, A to C). We then made a series of 100 nt deletions tiled across the segment from the Cap to nucleotide 739 and compared GFP expression from these reporters by immunoblot and fluorescence microscopy (Fig. 1, A to C, and fig. S1, A and B). RT-PCR assays of the expressed RNA showed equal RNA expression from each of the plasmids indicating that differences in GFP levels between constructs were not due to changes in the RNA levels (fig. S1, C and D). Deletions of *Malat1* nucleotides 640-739 and 540-639 greatly reduced GFP expression, while deletion of nucleotides 440-539 nearly eliminated it (Fig. 1, A to C). In contrast, deletion of nucleotides 1-139, 240-339, or 340-439 led to increased GFP expression.

**Fig. 1.**
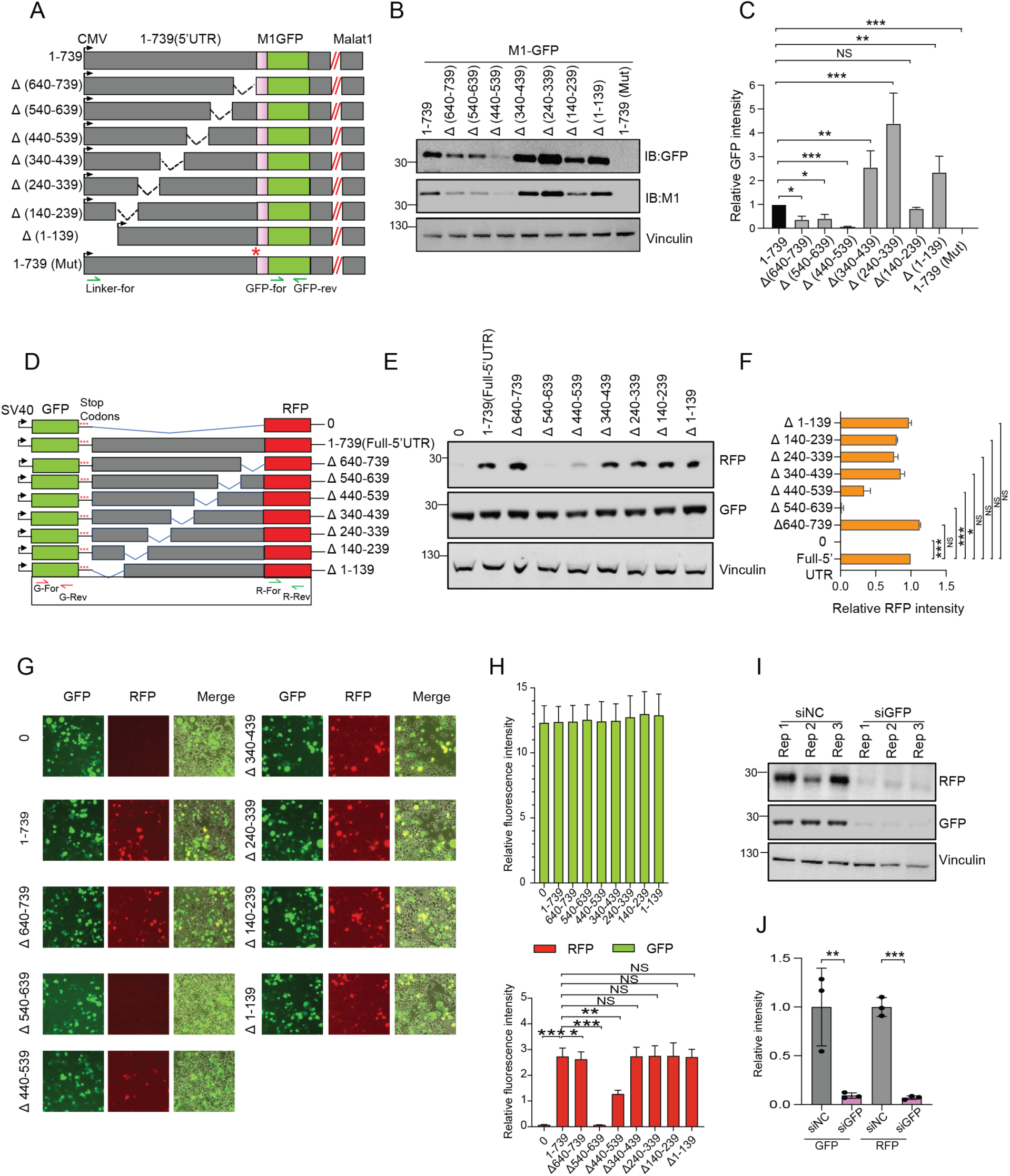
Identification of functional RNA segments required for M1 microprotein translation. (A) Diagram of constructs with GFP inserted in frame with M1 within the full-length Malat1 transcript, showing a series of 100-nucleotide deletions. The construct 1-739(Mut) contains a point mutation changing the M1 AUG to UAG (*). GFP-specific primer pairs and linker primer are indicated. (B) GFP and M1 immunoblot analysis of M1-GFP expression in N2a cells expressing the deletion constructs. Vinculin serves as a loading control. (C) Quantification of immunoblot results from (B). Bars represent mean ± SD; N = 3. (D) Diagram of GFP-RFP bicistronic reporters with the Malat1 5′ UTR (nucleotides 1-739) or 100-nucleotide deletions inserted between the two ORFs. GFP-and RFP-specific primer pairs are depicted. (E) Immunoblot analysis of GFP and RFP expression in N2a cells transfected with deletion constructs. Vinculin serves as a loading control. (F) Quantification of relative RFP intensity from (E), normalized to GFP. N = 3. (G) Fluorescence microscopy of GFP and RFP expression in N2a cells transfected with bicistronic deletion constructs. (H) Quantification of fluorescence intensity from (G). For each condition, 10 randomly selected fields were acquired and mean RFP intensity was normalized to mean GFP intensity. N = 3. (I) Immunoblot analysis of GFP and RFP in N2a cells transfected with the bicistronic reporter (1–739) and siNC or siGFP. (J) Quantification of immunoblot results from (I). ****, p < 0.0001; ***, p < 0.001; **, p < 0.01; *, p < 0.05; NS, not significant by unpaired Student’s t-test.

Earlier studies identified sequences in Malat1 affecting its nuclear localization(*22*–*24*), although these nuclear retention sequences are not in the 5’ UTR affected by our mutations. To examine whether our deletions were altering the abundance of the RNA in the cytoplasm, we performed fluorescent in situ hybridization (FISH) with probes covering the GFP coding sequence. Transcripts carrying the complete *Malat1* 5’ UTR were largely nuclear with approximately 10% of the RNA present in the cytoplasm, similar to the endogenous RNA, which is 5-10% cytoplasmic in N2A cells (*6*). Deletions of nucleotides 140-239, 540-639, or 640-739 all yielded levels of cytoplasmic Malat1 RNA similar to the complete 5’ UTR. Several deletions altered localization of the reporter RNA, including 240-339 and 340-439 that increased the cytoplasmic RNA and 440-539 that reduced it (Fig. S1, E and F). The loss of GFP protein expression observed from the 440-539 deletion is thus likely due in part to reduced cytoplasmic RNA available for translation. An earlier study of mouse Malat1 found that in HC11 mammary epithelial cells, deletion of the B1 SINE element from the endogenous gene led to increased cell numbers exhibiting detectible cytoplasmic Malat1 RNA(*23*). When this element (interval 540-639) was deleted from our reporter gene, we did not observe increased cytoplasmic RNA in N2A neuroblastoma cells. These different observations could arise from the different cell lines examined, as we previously found that endogenous cytoplasmic Malat1 is more abundant in neurons and N2A cells than in nonneuronal cells. Overall, we identified several RNA segments upstream of the M1 ORF affecting Malat1 localization and translation. We focused on segments 540-639 and 640-739 whose deletion in our system did not affect cytoplasmic RNA levels but reduced protein expression, and are apparently required for its translation.

We next tested whether the *Malat1* 5’ UTR affected M1 expression independently from the rest of *Malat1* RNA. The 739 nt UTR was placed between the GFP and RFP ORFs of a bicistronic reporter(*25*, *26*), with the RFP initiation codon in the same position as the M1 peptide (Fig. 1D and fig. S1G). The upstream GFP undergoes standard Cap dependent translation. Three stop codons were placed between GFP and the *Malat1* segment to prevent translational readthrough from GFP to RFP (fig. S1G). In the absence of the *Malat1* insert, no RFP expression is observed (Fig. 1, D to F). In contrast, the full *Malat1* 5’ UTR induces RFP expression, possibly indicative of an internal ribosome entry site stimulating RFP translation. We then tested the deletion series along the UTR for effects on RFP expression, assayed by both immunoblot and fluorescence microscopy (Fig. 1, D to H). All the clones expressed equal levels of GFP indicating that the mRNAs containing just the 5’ UTR fragment of Malat1 were not retained in the nucleus. In the context of this bicistronic reporter, RFP expression was again dependent on the portions of the Malat1 UTR present in the mRNA. Deletion of nucleotides 440-539 and 540-639 reduced expression of RFP (Fig 1, E and F), while deletions within the region from 1-439 had no effect (Fig. 1, E and F). With this reporter, loss of the final segment of the UTR between nucleotides 640-739 did not affect RFP expression (Fig. 1, E and F). Thus in both reporters, a segment of RNA that overlaps the B1 SINE strongly affected protein expression from the downstream ORF.

Assays of internal ribosome entry sites using bicistronic reporters are subject to several kinds of false positives(*27*, *28*). The presence of cryptic introns that remove upstream sequences or of cryptic promoters between the two ORFs that shorten the product mRNAs can allow cap dependent translation of the downstream ORF(*29–31*). To assess these possibilities, we measured RNA expression from the different reporters by RT-PCR. The full-length *Malat1* UTR construct and its mutants all showed constant expression of both the GFP coding sequence and of RNA extending from the linker upstream of the *Malat1* 5’ end to the GFP ORF (fig. S1, C and D). No shorter products were observed that might indicate removal of internal segments by splicing (fig. S1C). Similarly with the bicistronic reporter, the RNA segments individually encoding RFP and GFP as well as the long segment extending from GFP to RFP were expressed at constant levels by all clones (fig. S1, H and I). There was no evidence of splicing removing sequences upstream of RFP or of transcripts initiating within the *Malat1* insert (fig. S1H). To further confirm that the RFP expression was arising from RNA that contains the GFP ORF, we performed RNAi knockdown experiments. siRNAs targeting the GFP ORF eliminated both proteins, indicating that RFP was not coming from a shorter RNA initiating downstream of GFP (Fig. 1, I to J and fig. S1J). As described below, we also demonstrated that a minimal IRES sequence was independent of nuclear transcription and processing and independent of cap recognition. In sum, we found no evidence of alternative transcripts that might yield cap-dependent translation of RFP and conclude it is likely the product of IRES activity in the *Malat1* 5’ UTR.

### The mouse *Malat1* RNA sequence required for M1 translation forms a stable secondary structure *in vivo*

IRES elements function as stable, folded RNA structures that use a variety of mechanisms to recruit translation factors and ribosomal subunits to an internal segment of RNA (*32*, *33*). To assess the RNA folding of the 5’ region of *Malat1*, we performed four-base dimethyl sulfate mutational profiling (fbDMS-MaP) in mouse BV2 cells(*34–36*) (Fig. 2A). Cells were treated with DMS under conditions that allow the probing of all 4 nucleotides, the modified RNA was isolated, and a 1.1 kb segment of the *Malat1* RNA encompassing the 5’ end and the beginning of the M1 ORF was reverse transcribed. In this step, the DMS modifications introduce mutations into the cDNA. The cDNA was then amplified by PCR, sequenced, and mutation rates were analyzed by ShapeMapper 2.2 run in DMS mode(*36*). The individual base mutation rates for A, C, and U were highly dependent on the DMS modification (Fig. 2B and fig. S2A), suggesting *Malat1* was successfully modified. Following previous observations(*36*), the G base mutation rate was only moderately dependent on the DMS treatment (Fig. 2B and fig. S2A). DMS reactivities between the two biological replicates were highly reproducible with a Pearson correlation of 0.78 if all four bases were considered, and 0.93 for the reactivities of A, C, and U (Fig. 2C).

**Fig. 2.**
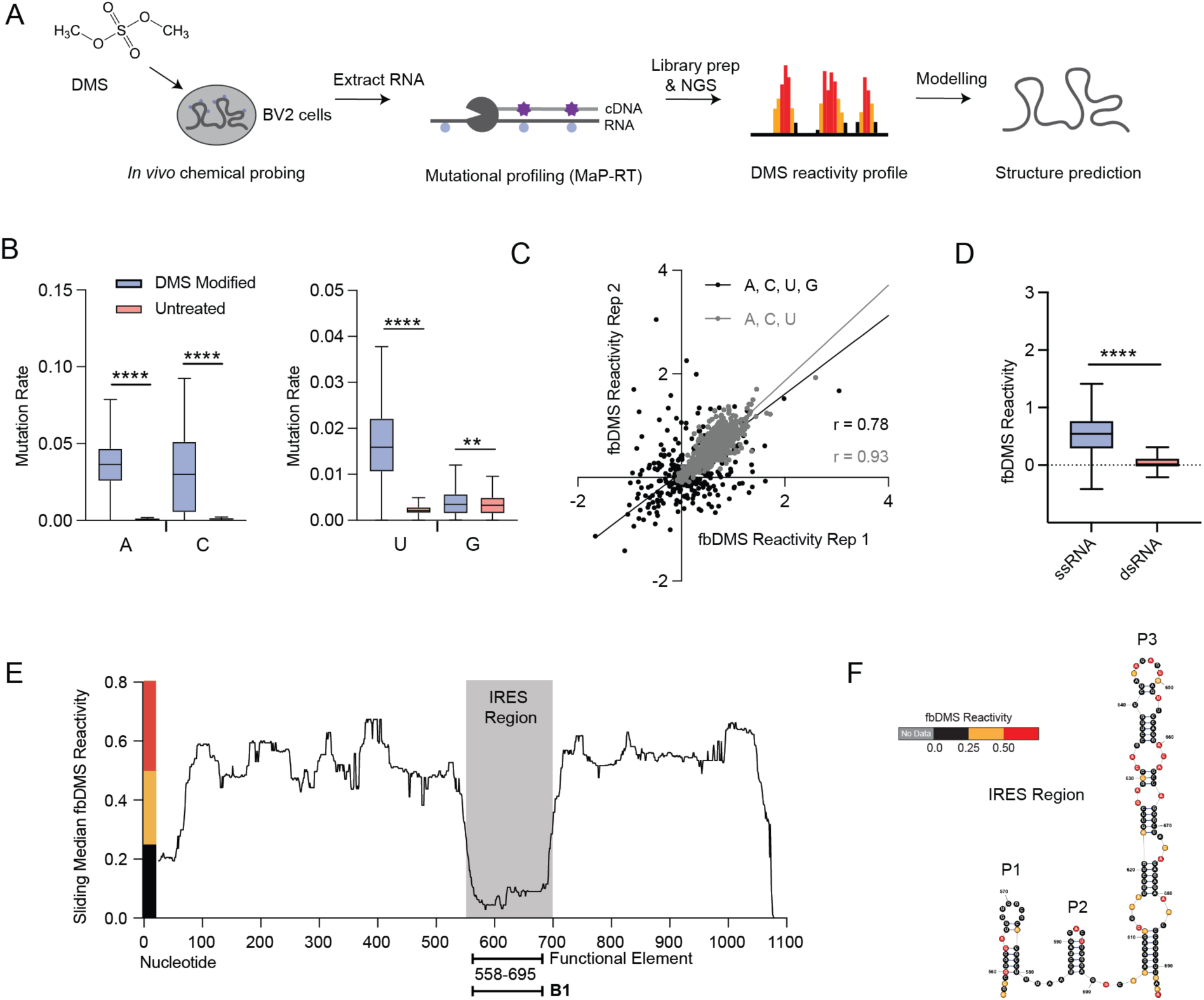
*In vivo* fbDMS probing reveals a defined RNA secondary structure within the 5’ UTR of endogenous mouse *Malat1*. (A) Chemical probing pipeline for *in vivo* four-base-DMS mutational profiling (fbDMS-MaP). (B) Mutation rates per nucleotide of one biological replicate of fbDMS-MaP probing for the 5’ region (1.1 kb) of mouse *Malat1* in BV2 cells. Boxes show the interquartile range, the median is indicated by a line, whiskers are drawn in Tukey style and values outside this range are not shown. (C) Correlation of fbDMS reactivities from two biological replicates of fbDMS-MaP probing of the 5’ region of mouse *Malat1* in BV2 cells. Each line is a linear regression fit to the data with corresponding Pearson correlation coefficients. (D) Mapping of fbDMS reactivities onto single-stranded (ssRNA) and double-stranded (dsRNA) nucleotides of the RNA secondary structure model of the mouse *Malat1* 5’ region. The model was predicted using SuperFold 1.2 with constraints from fbDMS reactivities. (E) Sliding median analysis of fbDMS reactivities along mouse *Malat1* 5’ region calculated in 51nt sliding windows. Regions of low (black), medium (orange) and high (red) fbDMS reactivity are shown. The regions containing the mouse *Malat1* IRES and SINE element are indicated. (F) RNA secondary structure model of the mouse *Malat1* IRES region colored by fbDMS reactivity. The three major stems are designated as P1, P2, P3. ****, p < 0.0001,***, p < 0.001; **, p < 0.01; *, p < 0.05 by equal variance unpaired Student’s t-test.

To identify structured segments in the *Malat1* sequence, we next performed secondary structure prediction using SuperFold 1.2 with the fbDMS data as experimental constraints. Within the consensus structure model generated by Superfold, the predicted double-stranded nucleotides exhibited significantly lower reactivities than predicted single-stranded regions, indicating good agreement between the fbDMS-MaP data and the secondary structure model (Fig. 2D and fig. S2B), although we can’t rule out that some regions of low reactivity stem from bound proteins. To identify regions exhibiting low reactivity, and hence potentially higher base-pairing content, we determined the median fbDMS reactivity within a 51 nucleotide sliding window across the 1.1kb *Malat1* RNA segment (Fig. 2E and fig. S2C). This reactivity profile identified an ∼150 nt interval of reduced reactivity between *Malat1* nucleotides 550 and 700 that both overlapped with segments whose deletion inhibited M1 translation and contained the B1 element. Superfold predicts a structure within this region consisting of two short hairpin loops P1 and P2, followed by a longer P3 structure of multiple base-paired stems separated by bulge loops and capped by a terminal loop (Fig. 2F and fig. S2D). The experimental reactivities map well onto the structure prediction, with single-stranded nucleotides exhibiting higher reactivies, while the base-paired stems contained lowly reactive nucleotides (Fig. 2, D to F, and fig. S2, B to D). Overall, the fbDMS-MaP data support the presence of a stable RNA structure within the sequence required for M1 translation.

We next performed more refined mutagenesis to assess the IRES activity of the structure defined by fbDMS-MaP and of the B1 element. Deletions upstream and downstream of the structure did not strongly affect RFP expression, except for a short segment immediately upstream (nucleotides 540-557), whose deletion showed moderate reduction in RFP (Fig. 3, A to D, and fig. S3, A-B). Deletion of either P1 or P3 severely reduced RFP translation, while deletion of P2 had a partial effect (Fig. 3, A to D, fig. S3B). Thus, the IRES activity within the *Malat1* 5’ region correlates well with the secondary structures defined within this RNA. To determine the minimal sequence needed for IRES activity, we created additional reporter constructs containing just the structured region or portions of it (Fig. 3E). The segment containing just the P1, P2, and P3 structures (nucleotides 558-695) was sufficient to induce RFP expression to the same level as the full 5’ UTR (Fig. 3, E-G). Adding nucleotides 540-557 upstream of P1 to this minimal IRES did not increase its activity(Fig. S3 I-J), but trimming its 3’ end to remove a portion of P3 (640-695) greatly reduced activity (Fig. 3, E-G). Smaller fragments containing just P1, P2, or P3, or the P1-P2 or P2-P3 segments all lacked IRES activity (Fig. 3, E-G). The minimal active segment (mini-IRES) notably coincided with both the defined structural elements (P1+P2+P3) and the annotated B1 SINE (nucleotides 556 to 703) (Fig.S2D).

**Fig. 3.**
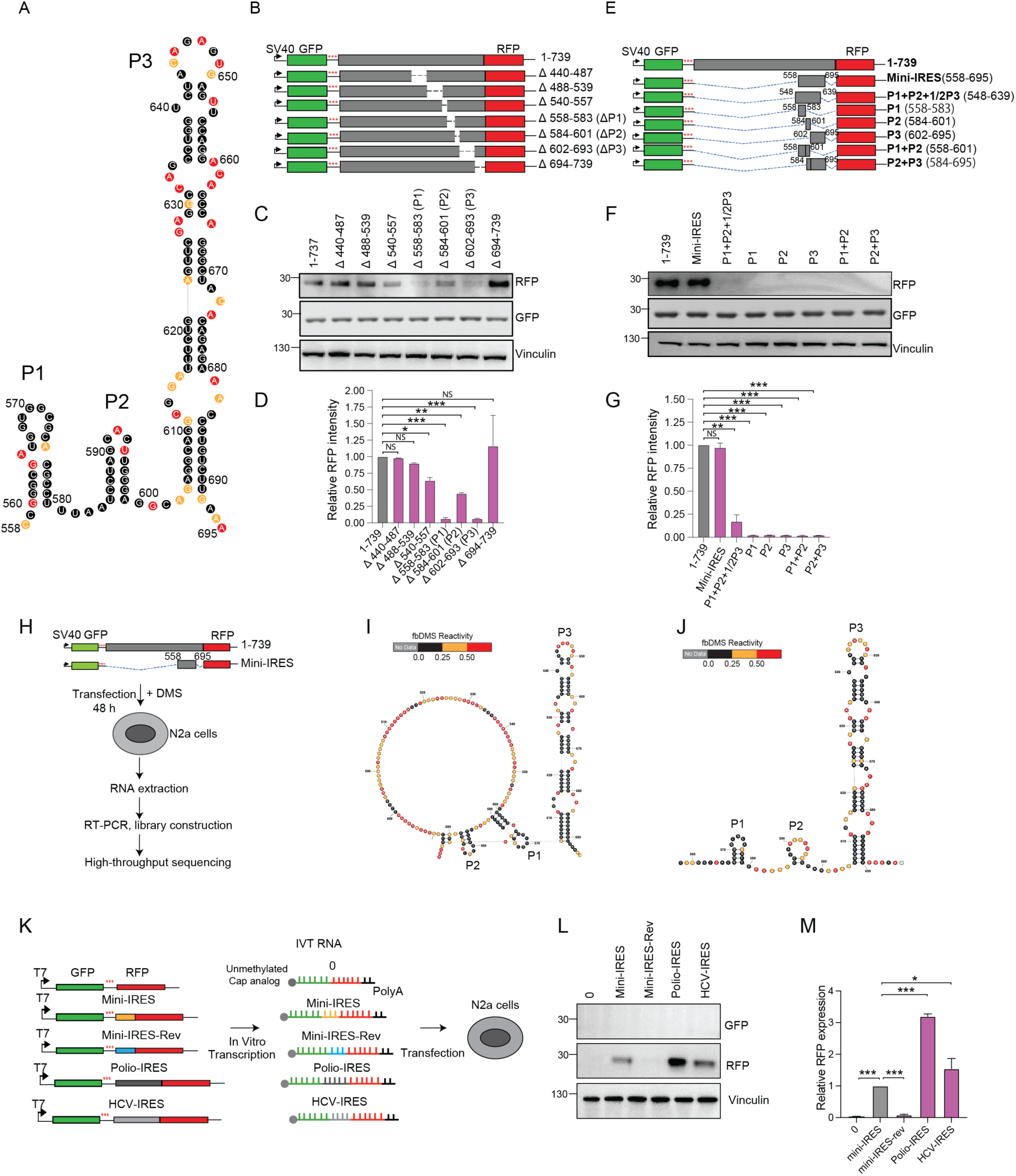
The secondary structure within the *Malat1* 5’ UTR is essential for *Malat1* IRES activity and is maintained in the reporter RNAs. (A) Enlarged secondary structure model of endogenous Malat1 (nucleotides 558–695) determined by fbDMS probing for comparison to deletion mutants. The three stems are designated P1, P2, P3. (B) Diagrams of GFP-RFP bicistronic reporters with the Malat1 5′ UTR positioned between the two cistrons, carrying deletions of predicted secondary structures. (C) Immunoblot analysis of RFP and GFP expression in N2a cells transfected with constructs from (B). (D) Quantification of relative RFP expression from (C). N = 3. (E) Diagrams of bicistronic constructs carrying different segments encompassing the secondary structure motifs inserted between the GFP and RFP ORFs. (F) Immunoblot analysis of RFP and GFP expression in N2a cells transfected with constructs from (E). (G) Quantification of relative RFP expression from (F). Bars represent mean ± SD; N = 3. (H) Pipeline for fbDMS probing of secondary structures within the Malat1 5′ UTR (1–739) and minimal IRES reporter constructs. (I) RNA secondary structure model of the Malat1 5′ UTR (1–739 nt) reporter colored by fbDMS reactivity. (J) RNA secondary structure model of the Malat1 minimal IRES (558–695 nt) reporter colored by fbDMS reactivity. (K) Diagram of in vitro transcribed bicistronic RNAs with unmethylated cap analog and poly(A) tail transfected into N2a cells. (L) Immunoblot analysis of RFP and GFP expression in N2a cells transfected with in vitro transcribed RNAs. HCV and Polio IRES serve as positive controls; no insertion and reversed IRES serve as negative controls. (M) Quantification of relative RFP expression from (L). Bars represent mean ± SD; N = 3. ****, p < 0.0001; ***, p < 0.001; **, p < 0.01; *, p < 0.05 by unpaired Student’s t-test.

The initial fbDMS-MaP was performed on full-length endogenous *Malat1*, which may adopt a different structural conformation compared to our reporter RNAs. To assess the structure of the IRES region within the RNA expressed from the reporter gene, we repeated the fbDMS-MaP on RNA from mouse N2A cells transfected with plasmids containing the full 739 nucleotides from the *Malat1* 5’ UTR or containing just the minimal IRES (nucleotides 558-695; Fig. 3H). Cells transfected with the respective plasmids were treated with DMS and processed as described above. In these experiments, the mutation rates in the modified samples were significantly higher than the untreated samples for all four bases (fig. S3, C and F). The reactivities of the biological replicates were highly correlated for all four bases with r = 0.96 for the full-length construct and r = 0.99 for the minimal IRES (fig. S3, D and G). As before, the fbDMS reactivities were significantly decreased for the predicted double-stranded nucleotides compared to single-stranded nucleotides within the structures computed by Superfold (fig. S3, E and H). The consensus structure model for the full-length UTR was almost identical to the endogenous RNA (Fig. 3A and I), while the minimal IRES also formed P1, P2, and P3 but showed the lower nucleotides of the P1 and P2 stems as unpaired (Fig. 3, A and J). Overall the fbDMS-MaP data indicate that the IRES RNA likely folds into a similar structure in all three sequence contexts. This predicted secondary structure of the Malat1 IRES in vivo is in close agreement with a B1 SINE RNA structure determined by phylogenetic comparison and enzymatic probing in solution (*37–40*), with P1, P2 and P3 segments all present(Fig S4, A-C).

To further confirm the activity of the minimal IRES and to compare it to other IRES’s, we created RNAs by *in vitro* transcription. Bicistronic RNAs were synthesized with different sequences separating the ORFs, including no insert, the minimal *Malat1* IRES, the inverted sequence of the minimal IRES, or the well characterized IRES’s from Poliovirus and Hepatitis C Virus (HCV) (Fig. 3K and fig. S4H). The RNAs were capped with unmethylated G and terminated in a polyA sequence encoded by the runoff template (Fig. 3K). The unmethylated cap is not expected to support cap dependent translation of GFP but should not affect IRES dependent translation of RFP. These RNAs were transfected into N2a cells and assayed for GFP and RFP expression by immunoblot (Fig. 3K). As expected, none of the RNAs yielded GFP expression. The RNAs with no insert or with the reversed IRES produced no RFP. In contrast, the minimal Malat1 IRES, the polio IRES, and the HCV IRES all expressed RFP. The *Malat1* and HCV IRES exhibited about 1/3 the activity of the Polio IRES, with HCV slightly more active than Malat1 (Fig. 3, K to M). IRES elements are known to vary in their required cellular cofactors and these differences in activity may reflect the relative abundance of these factors in the N2A cells. To confirm the cap dependence of the GFP translation and assess how this might affect the RFP ORF, we repeated the experiment with three RNAs carrying 7-methyl Guanosine caps (fig. S4, D-E). The methylated cap supported GFP translation but did not strongly affect translation of RFP, with the Malat1 IRES again exhibiting about 1/3 the activity of the polio IRES (fig. S4, F-G). These data confirm that the *Malat1* IRES functions without the need for nuclear transcription and processing.

### The mouse *Malat1* IRES binds to ribosomal subunits and translation factors

IRES elements recruit ribosomes to an mRNA without the need for cap-dependent translation initation, but differ in their mechanisms of recruitment and required cofactors(*32*). To identify factors that bind the *Malat1* IRES we performed RNA affinity pulldown experiments. The minimal IRES was linked to three copies of the streptavidin binding RNA aptamer (3xS1m), transcribed *in vitro*, and immobilized on Streptavidin Sepharose beads (Fig. 4A and fig. S5A)(*41*, *42*). As a length-matched control, the reverse sequence of the minimal IRES was also fused to the streptavidin aptamer. We also created and immobilized IRES RNAs that were lacking either the P1 or the P3 stem loops, or contained just the streptavidin aptamer (Fig. 4A and fig. S5A). The beads were incubated in lysates of N2a cells, washed, and the RNA-bound factors were eluted with ribonuclease and displayed by SDS-PAGE (fig. S5B). The gel profiles showed some proteins present in all the eluates including the control reverse IRES and the streptavidin aptamer alone (fig. S5B). Along with these nonspecific binders, the full length IRES bound to an array of additional proteins. The IRES elements lacking P1 and P3 bound subsets of the proteins seen with the full IRES (fig. S5B and table S1).

**Fig. 4.**
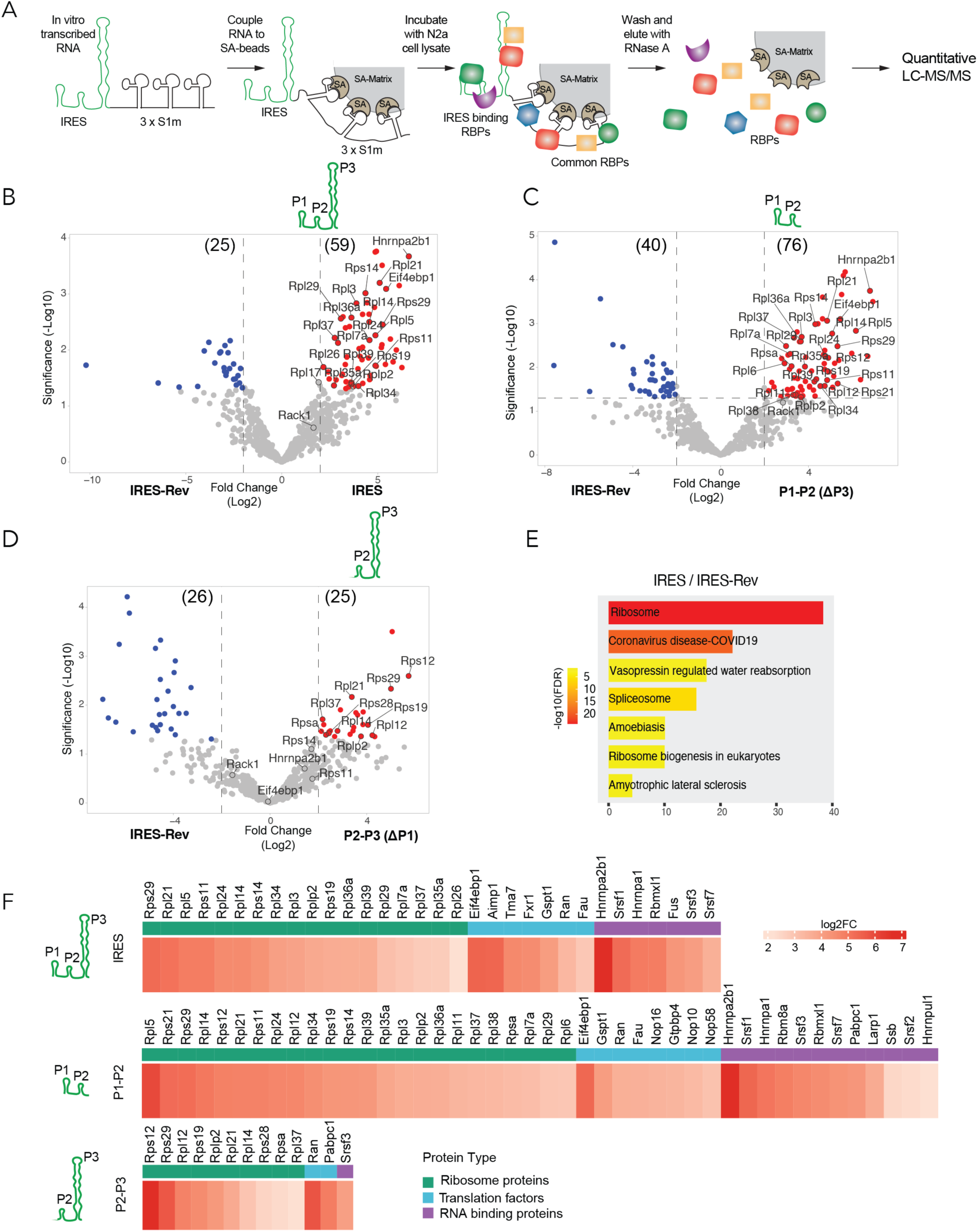
Identification of IRES binding proteins. (A) Schematic of the S1m-IRES RNA affinity pulldown assay. (B) Volcano plot of protein abundances determined by MS in the IRES eluate compared to the IRES-rev. Proteins with a fold-change greater than 4 and P values less than 0.05 are colored red. Proteins with a fold change less than 0.25 and a p-value < 0.05 are shown in blue. Proteins with intermediate values are shown in grey. (C) Volcano plot of protein abundances determined by MS in the IRES-ΔP3 eluate compared to the IRES-rev. Proteins with a fold-change greater than 4 and P values less than 0.05 are colored red. (D) Volcano plot of protein abundances determined by MS in the IRES-ΔP1 eluate compared to the IRES-rev. Proteins with a fold-change greater than 4 and P values less than 0.05 are colored red. (E) GO analysis of proteins significantly enriched in the IRES eluate compared to IRES-rev. (F) Heatmap illustrating the variable enrichment of ribosomal proteins, translation factors, and RNA-binding proteins associated with the mini-IRES, P1/P2, and P2/P3 RNAs.

Replicate samples from the five different RNA pulldown conditions were characterized by LC-MS/MS mass spectrometry using label free quantification(*43*). This identified 617 proteins isolated across the five conditions (table S1). Proteins were ranked by abundance in the IRES sample, with the top 300 used for comparison to the other pulldown conditions. Proteins across the five samples were clustered by relative abundance and colored using row-wise scaling (fig. S5C). This displayed clusters of proteins with strong preferential binding to the full IRES or to both the full IRES and the IRES with P3 deleted (P1-P2). Smaller groups of proteins preferentially bound the negative control RNAs containing S1m alone or the reverse IRES sequence, as well as the IRES with P1 deleted (P2-P3) (fig. S5C). The proteins specifically binding to the IRES included many ribosomal proteins and translation factors. They also included SRP9 and SRP14 that are expected to bind the B1 element, although these were less abundant than the ribosomal proteins and others (Table S1). Some proteins involved in splicing also bound to the IRES, including hnRNPA2B1, hnRNPA1, Rbmxl1, Srsf1 and Srsf7, as well proteins annotated for different functions (Fig. 4F and table S1). Histone proteins bound both the full IRES and the reverse IRES over the other shorter RNAs, indicating their affinity may depend on the length of tested RNA (fig. S5C). Examining the IRES-bound proteins by gene ontology for enriched cellular components and biological processes identified ribosome-related terms as the most significant functional category in both the IRES and P1-P2 samples (Fig. 4E and fig. S5E). Proteins with preferential binding to the reverse IRES or the Streptavidin tag alone were enriched for other non-ribosomal categories (fig. S5D, table S1).

Proteins binding the full IRES, P1-P2 and P2-P3 RNAs were displayed in volcano plots according to their fold enrichment over the reverse RNA control. Using cutoffs of a 4-fold enrichment over the control eluate and a p value of less than 0.05, we identified 59 proteins that specifically associate with the full IRES, 76 with the P1-P2 (P3 deletion), and 25 with the P2-P3 (P1 deletion) over the reverse IRES control. (Fig. 4, B to D and table S1). The presence of both large and small ribosomal subunit proteins suggests binding of 80S ribosomes. However, not all ribosomal proteins were equally represented in the bound fractions (Fig. 4F), possibly due to fragmentation of the ribosome during the RNase elution or selective loss during the wash step before elution. Deletion of P1, which eliminated IRES activity *in vivo*, caused loss of binding for multiple proteins including hnRNP A2/B1, Rps11, Rps14, and Rack1, as well as the B1 binders SRP9 and SRP14 (Fig. 4, C to F). Overall the affinity purification data confirm that ribosomal subunits are binding to the *Malat1* IRES (*29*, *42*, *44–48*).

### IRES binding proteins Rack1 and hnRNPA2/B1 are required for *Malat1* IRES activity

We confirmed selected IRES binding proteins by immunoblot. Rps14, Rps11, hnRNP A2/B1, and Rack1 all exhibited substantial binding to the IRES and P1-P2 RNAs but were present in only trace amounts in the 3xS1m, 3xS1m-Rev, and P2-P3 eluates (Fig. 5A). Similar to the MS results, the protein Rack1 was moderately enriched in the IRES eluate, and this binding was lost after deletion of the P1 stem loop but enhanced by deletion of P3 (Figs. 4, B to D and 5A). Rack1 is a component of the small ribosomal subunit that is not required for Cap dependent translation initiation but is essential to the activity of several viral IRES elements(*49–52*). To test its effect on the *Malat1* IRES, we carried out siRNA mediated depletion of Rack1 in cells expressing the full-length or the minimal IRES reporters. We also tested the effect of increasing Rack1 levels by transgene expression. Assayed by immunoblot, siRNA treatment led to an ∼70-80% depletion of Rack1 protein from the N2A cells, without altering expression of other small ribosomal subunit proteins (fig. S6G). For both the full-length and the minimal IRES reporters, this led to ∼75% decrease in RFP expression from the reporter, without affecting expression of the upstream GFP (Fig. 5, B and C). Conversely, overexpression of Rack1 boosted RFP expression about 3-fold, again with no change in GFP (Fig. 5, D to E). Similarly, when assayed by fluorescence microscopy, the pixel intensity of Rack1 from the transfected plasmid showed strong positive correlation with RFP fluorescence produced from the mini-IRES reporter (r=0.823; fig. S6, A and B). The translation induced by the *Malat1* IRES is thus dependent on Rack1.

**Fig. 5.**
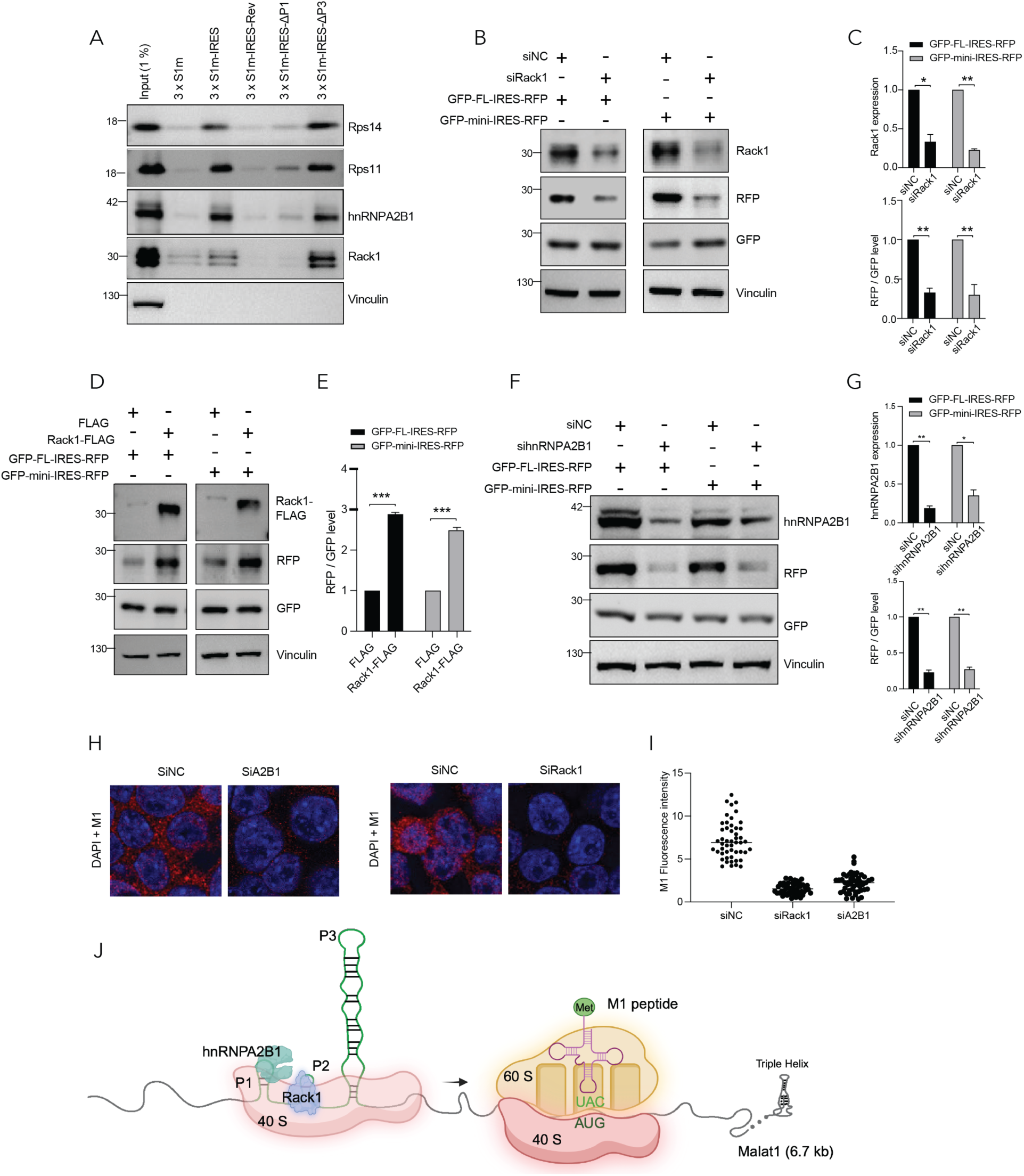
*Malat1* IRES activity is dependent on Rack1 and hnRNP A2/B1. (A) Immunoblot analysis of selected proteins enriched in the IRES eluate compared to S1m, IRES-rev, IRES-ΔP1, and IRES-ΔP3 eluates. (B) Immunoblot analysis of RFP, GFP, and Rack1 in N2a cells transfected with control or Rack1 siRNAs and the GFP-FL-IRES(1-739)-RFP (left) or GFP-mini-IRES(558-695)-RFP (right) plasmid. Vinculin serves as a loading control. (C) Top: quantification of Rack1 knockdown efficiency from (B). Bottom: quantification of relative RFP expression normalized to GFP from (B). Bars represent mean ± SD; N = 3. (D) Immunoblot analysis of RFP, GFP, and Flag-Rack1 in N2a cells transfected with GFP-Flag or Flag-Rack1 together with GFP-FL-IRES-RFP (left) or GFP-mini-IRES-RFP (right). Vinculin serves as a loading control. (E) Quantification of relative RFP expression normalized to GFP from (D). Bars represent mean ± SD; N = 3. (F) Immunoblot analysis of RFP, GFP, and hnRNP A2/B1 in N2a cells transfected with control or hnRNP A2/B1 siRNAs and GFP-FL-IRES-RFP(1-739) or GFP-mini-IRES-RFP(558-695) plasmids. (G) Top: quantification of hnRNP A2/B1 knockdown efficiency from (F). Bottom: quantification of relative RFP expression normalized to GFP from (F). Bars represent mean ± SD; N = 3. (H) Fluorescence imaging of M1 expression in N2a cells transfected with siNC, siA2B1, or siRack1. (I) Quantification of M1 fluorescence intensity in single cells transfected with siNC, siA2B1, or siRack1. N = 50. (J) Diagram of proposed ribosome recruitment to the Malat1 IRES. ****, p < 0.0001; ***, p < 0.001; **, p < 0.01; *, p < 0.05 by unpaired Student’s t-test.

The most enriched protein in the IRES and P1-P2 samples was hnRNP A2/B1 (Fig. 4, B to D and F). This protein is known to act as an internal ribosome entry site trans-acting factor (ITAF) that promotes translation of several viral and cellular mRNAs, including Glutamate Receptor A1 mRNA whose translation is localized to neuronal dendrites similar to *Malat1*(*53–56*). To verify its role in the *Malat1* IRES, siRNA knockdown was performed in cells expressing reporter genes. In cells expressing either the full 5’ UTR or the minimal-IRES reporter hnRNP A2/B1 expression was reduced to 25% and 40% that of control cells, respectively (Fig. 5, F and G). Similar to Rack1 depletion, immunoblot and fluorescence assays showed that hnRNP A2/B1 depletion significantly reduced RFP expression without changes in GFP expression (Fig. 5, F and G and S5, C to F).

To confirm that M1 peptide expression from the endogenous MALAT1 RNA was also dependent on Rack1 and hnRNP A2/B1, we assayed M1 immunofluorescence after knockdown of the two proteins. Loss of either Rack1 or A2/B1 led to a dramatic reduction in M1 peptide fluorescence without changing expression of other small ribosomal subunit proteins (Fig. 5, H to I, fig.S6, G to H). Thus, like the RFP protein produced from reporter genes, endogenous M1 peptide expression requires the IRES cofactors. M1 peptide translation from the endogenous Malat1 transcript is thus likely mediated by the IRES (Fig. 5, H-J).

## Discussion

We identify a B1 SINE element that acts as internal ribosome entry site required for the translation of a micropeptide encoded in the mouse *Malat1* RNA. IRES elements are diverse in structure and cofactor dependencies and can employ different mechanisms of ribosome recruitment (*29*, *32*, *57*, *58*). IRES’s of encephalomyocarditis virus (EMCV) and poliovirus recruit the eIF4G/eIF4A complex to initiate ribosome assembly(*46*, *59–63*), whereas the IRES in human Hepatitis C virus recruits eIF3, Rack1, and other factors. Other IRES’s may recruit ribosomal subunits via direct RNA interactions. We are not aware of previous reports of B1 elements having IRES activity, but human Alu SINE elements are known to bind 80S ribosomes(*64*). A screen for IRES elements across thousands of human cellular and viral RNA segments identified a large set of candidate IRES’s that grouped into several classes(*58*). Among these we find several that contain Alu elements, although it is not yet clear whether the SINE is important for their activity.

An earlier study examined a deletion of the B1 element from the endogenous Malat1 of mouse HC11 mammary epithelial cells(*23*). In this study, deletion of Malat1 caused an increase in the number of cells showing cytoplasmic Malat1 RNA. This was tied to an increase in cytoplasmic TDP43 protein and several physiological defects. We did not observe increased cytoplasmic RNA upon deletion of the B1 element. Instead, we identified sequences adjacent to the IRES that affected the nucleo-cytoplasmic distribution of the RNA. This discrepancy could be due to the different cell lines used, different behaviors of the endogenous and plasmid-expressed RNAs, the different ways that cytoplasmic RNA was quantified, or some other factors. N2A neuroblastoma cells, like neurons, express more cytoplasmic RNA than non-neuronal cells we have tested. It is possible that factors affecting the Malat1 nucleo-cytoplasmic ratio vary between cells.

B1 SINEs are abundant sequence repeats found throughout the mouse genome that are analogous to human Alu elements. Cytoplasmic MALAT1 and MALAT1-derived peptides have been observed in human cells(*65*, *66*). However, B1 elements are specific to rodent genomes and the human MALAT1 RNA does not contain an Alu or similar element adjacent to its putative M1 ORF. Additional studies will be needed to assess the mechanisms leading to translation of human MALAT1. fbDMS-MaP analyses of endogenous *Malat1* RNA in mouse, and of reporter gene transcripts containing the IRES, all predicted structures consistent with that of the B1 element (*39*, *40*) (Fig. 3, A, I and J). The fbDMS-MaP model consists of two short stem loops P1 and P2 upstream of a more extended P3 stem containing several bulge loops. These features are all present in the B1 structure(fig S4, B-C). An additional H0 helix in the SINE structure that pairs a short segment upstream of P1 to nucleotides preceding P3 is also consistent with the Malat1 probing data (fig S4, B-C). In a solved 3-dimensional structure of the human Alu RNP, P1 and P2 with their terminal loops and connecting linker form a stable fold that docks against the P3 stem(*38*, *64*). This Alu RNA fold is consistent with observed structure of the mouse Malat1 IRES.

B1 and Alu elements are both derived from the 7SL RNA(*40*, *67*, *68*), a component of the signal recognition particle (SRP) that arrests translation of secreted proteins prior to their recruitment to the ER membrane(*69*, *70*). This translation arrest occurs by SRP blocking the eEF2 binding site at the interface of the two ribosomal subunits(*71*, *72*). B1 and Alu differ from 7SL in having acquired features that enhance their transposition by LINE element proteins. The B1 element RNA structure retains binding of the SRP9 and SRP14 proteins, while Alu elements are a dimer of this structure(*73*, *74*). Interestingly, the Alu RNP causes translational arrest similar to SRP and this is thought to allow translating LINE RNPs to act on SINE RNAs during transposition(*64*, *75*, *76*). It is interesting to speculate that similar interactions of the Malat1 B1 element may serve to recruit ribosomes for translation of the M1 peptide.

In affinity purification experiments, the most prominent group of IRES binding proteins were ribosomal proteins, translation factors, and other RNA binding proteins. Bound proteins included components of both the small 40S and large 60S ribosomal subunits (Rps and Rpl proteins) (Fig. 4 and table S1) indicating likely binding of full 80S ribosomes. It is not clear if these are directly recruited, perhaps similarly to SRP, or if there is initial recruitment of the small subunit as seen with most viral IRES. Determining the mechanism of Malat1 IRES function and comparing it to other IRES like HCV will require studies using *in vitro* translation systems.

Many proteins bound to the full IRES also bound to the truncated P1-P2 RNA (lacking P3), possibly indicating that P1-P2 is sufficient for some ribosome interactions (Fig. 4, B-D and fig. S5B). Among the P1 dependent proteins was Rack1 (Figs. 4, B to F, and 5A). Rack1 is a small subunit ribosomal protein that is not essential for general translation but plays an important role in the cellular response to stalled ribosomes and resolving ribosome collisions(*77*). Rack1 is essential for the activity of several viral IRESs including HCV (*49*). We find that Rack1 strongly promotes IRES dependent RFP translation but has no effect on cap-dependent GFP expression. It thus appears to be key factor in the function of the *Malat1* IRES.

Another protein specifically binding to the IRES was hnRNP A2/B1. Like Rack1, depletion of hnRNP A2/B1 inhibits IRES dependent translation of the M1 peptide from the endogenous Malat1 RNA(Fig. 5H–5I). HnRNP A2/B1 is also required for IRES dependent translation of the Glutamate Receptor GluA1 transcript (Gria1), and of Senecavirus(*54*),(*53*). The GluA1 mRNA has several interesting similarities to *Malat1*. GluA1 mRNA is transported to dendrites where it is locally translated. GluA1 translation in Hippocampal neurons is stimulated by the neurotrophic factor BDNF, while *Malat1* translation in cortical cultures is stimulated by cellular depolarization with KCl. It will be interesting to examine whether *Malat1* IRES activity is responsive to neuronal stimulation. HnRNP A2/B1 is also reported to mediate the transport of mRNAs into cellular processes similar to the transport of *Malat1* in neurons(*78–80*).

Other proteins bound by the Malat1 IRES have been identified as IRES transacting factors (ITAFs), including hnRNP A1, eIF4ebp1, and Srsf3. hnRNP A1 binds the 5’ untranslated regions (UTR) of multiple mRNAs and viral RNAs to facilitate their translation(*81–83*). EIF4ebp1 (4E-BP1) is an inhibitor of cap-dependent translation through its sequestration of eIF4e, and EIF4ebp1 overexpression was found to promote IRES-dependent translation after cellular stress(*84*). SRSF3 facilitates IRES-dependent translation of Picornaviruses through interactions with the ITAF PCBP2(*85*). Determining the relevance of these many factors and identifying their interactions on the Malat1 RNA will require extensive mutagenesis studies and development of in vitro assays for IRES activity. Further dissection of the structure and factor dependencies of the *Malat1* IRES will allow its comparison with other viral and cellular IRESs and begin to resolve its mechanism of action. Of particular interest is the role of SRP9/SRP14 in ribosome binding. These proteins are seen to bind the IRES in the affinity purification but they are substoichiometric relative to the ribosomal proteins and their significance is not yet clear.

*Malat1* is an unusual RNA. Its novel 3’ end triple helix structure was previously shown to support translation despite the lack of a polyA tail(*18*). We now define a functional structure near the 5’ end of the mouse RNA that is required for translation of the M1 micropeptide. In the reporter RNAs, this IRES does not require the Malat1 3’ terminal triple helix to induce translation. However, translation of the M1 peptide from the endogenous Malat1 requires the IRES cofactors Rack1 and hnRNP A2/B1. The endogenous RNA thus has structures that allow its translation without either of the 5’ or 3’ features typically required by mRNAs. ORFs in more downstream portions of *Malat1* RNA may also be translated and there may be additional undescribed functional features within this interesting long RNA.

## Materials and Methods

### Cell lines

N2A and HEK293 cells were cultured in DMEM with 10% fetal bovine serum (FBS) and 10 U/ml Penicillin-Streptomycin. BV2 cells(*86*) were cultured in DMEM with 5% FBS, 1% non-essential amino acid solution and 10 U/mL Penicillin-Streptomycin. Cells were cultured in a CO2 incubator at 37 °C.

### Western Blotting

Cultured cells were lysed in RIPA buffer (150 mM sodium chloride, 50 mM Tris-HCl, 1 mM EDTA, 1% nonidet P40, 0.1% SDS, 0.5% Sodium deoxycholate) with protease and phosphatase inhibitors (Roche, 5056489001) and benzonase (Sigma Aldrich, E1014-25KU). Lysates were centrifuged, cleared, and boiled at 95°C for 5 min in NuPAGE 1x LDS sample buffer (Thermo, NP0007). Samples were loaded on a 4-12% Bis-Tris SDS-PAGE Protein Gels (NP0335BOX) and resolved in MOPS buffer (Thermo, NP0001). Protein Gel was transferred to 0.45 mm PVDF membranes (GE Amersham) or NC membrane (FisherScientific, PI88018) using semi-dry transfer machine (Biorad, 1703940). After 0.5 h blocking with 3% milk in PBST (1XPBS + 0.1% tween-20), the membrane was incubated with primary antibodies against protein of interest overnight. Membranes were washed 3 times with PBST and secondary antibody was added for 1 h incubation. After 3 times wash with PBST, the membranes were imaged on an iBright FL1500 Imaging System.

### Protein isolation on Streptavidin aptamer beads

1. Cell harvest: 3 x 15 cm dishes of cells were harvested with trypsin, transferred to 2 ml round bottom tubes (Eppendorf, 022363352), and snap-frozen in liquid nitrogen and stored at -80 for long-term use.
2. Homogenization of cells: A steel bead (diameter: 5-mm) was added into each 2 ml tube for homogenization, and cells were then homogenized on a Qiagen TissueLyser III twice at 25 Hz for 2 min.
3. Cell lysis: Homogenized cells were lysed by addition of 400 µL cold SA-RNP lysis buffer (20 mM Tris-HCl PH7.5, 150 mM NaCl, 1.5 mM MaCl_2_, 2 mM DTT, and EDTA free-protease inhibitors) for 10 min on ice with pipetting up and down. Cell debris was removed by centrifugation for 10 min at 17,000 x g at 4°C, resulting in a supernatant of ∼400 µL. This centrifugation clearance step was repeated 3 times to remove cell debris and lipids. The protein concentration was measured by Nanodrop.
4. Pre-clearing: 50 µL of 50% slurry of streptavidin Sepharose High Performance (GE healthcare) beads were washed 3 times with 1 ml of SA-RNP lysis buffer. The cell lysate was pre-cleared by addition of 50 µL streptavidin (SA) sepharose beads and tumbled for 0.5 h at 4C. Beads were discarded, the pre-cleared lysate was supplemented with 5 µL of RNase OUT inhibitor.
5. RNA-coupling with SA sepharose beads: 100 µL of a 50% slurry of SA sepharose beads (GE healthcare) were washed 2 times with 1 ml of SA-RNP lysis buffer. Beads were gently pelleted at 1000 rpm (∼ 100 xg) for 1 min at RT. 25 ug of *in vitro* transcribed IRES, IRES-rev, P1+P2(ΔP3), P2+P3(ΔP1) and 3xS1m RNAs were renatured in 50 µL SA-RNP buffer (supplemented with 0.5 µL RNaseOUT) by heating at 56°C for 12 min, then 10 min at 37°C, and then incubation at room temperature for 5 min to refold RNA structures. The 50 µL refolded RNA in SA-RNP buffer was added to washed SA-sepharose beads supplemented with 100 µL SA-RNP lysis buffer and incubated 0.5 h at 4°C under rotation. After centrifugation, the unbound RNA in the supernatant was removed, and the pelleted beads were kept for next step.
6. Protein binding to the RNA beads: The pre-cleared cell lysate was added to RNA-coupled SA sepharose matrix and incubated at 4°C overnight under rotation.
7. Washing and elution: Beads were washed 5 times (5 min each) with 1 ml SA-RNP Wash buffer (20 mM Tris-HCl PH7.5, 500 mM NaCl, 5 mM MaCl_2_, 2 mM DTT, and EDTA free-protease inhibitors). After the last wash, the beads were eluted by addition of 4 ng/ µL RNase A (thermo, AM2271) in 50 µL Low salt buffer (20 mM Tris-HCl PH7.5, 30 mM NaCl, 5 mM MaCl_2_, 2 mM DTT) for 10 min at 4°C. The eluates were analyzed by mass spectrometry, silver staining, and Western blotting.

### Liquid Chromatography-Tandem Mass Spectrometry (LC-MS/MS)

Eluates obtained from RNA pulldown were reduced with 5 mM tris(2-carboxyethyl)phosphine and subsequently alkylated with 10 mM iodoacetamide. The reduced and alkylated proteins were purified using the protein aggregation capture (PAC) method(*87*), followed by overnight proteolytic digestion at 37 °C with Lys-C and trypsin. The resulting peptides were dried prior to LC-MS/MS analysis.

Dried tryptic peptides were resuspended in 5% formic acid and injected onto a PepSep C18 reverse-phase column (150 mm × 150 µm, 1.7 µm particle size) maintained at 59 °C, followed by electrospray ionization into an Orbitrap Astral mass spectrometer. Chromatographic separation was performed using a trap-and-elute workflow on a Vanquish Neo UHPLC system. Mobile phase A consisted of water containing 0.1% formic acid, and mobile phase B consisted of acetonitrile with 0.1% formic acid. A 15-minute gradient was applied as follows: 0–1 min, 5% B (2.45 µL/min); 1–5 min, 5–15% B (1.75 µL/min); 5–12.6 min, 15–25% B (1.75 µL/min); 12.6–13.6 min, 25–38% B (1.75 µL/min); 13.6–13.7 min, 38–80% B (2.45 µL/min); and 13.7–15 min, 80% B (2.45 µL/min).

Data-independent acquisition (DIA) was carried out on the Astral mass spectrometer in positive electrospray ionization mode. MS1 spectra were acquired at a resolution of 240,000 over an m/z range of 380–980, with a normalized AGC target of 500% and a maximum injection time of 3 ms. DIA was performed using sequential 4 m/z-wide isolation windows across the 380–980 m/z range. MS2 spectra were acquired at a resolution of 80,000, employing higher-energy collisional dissociation (HCD) at a normalized collision energy of 25%, with a normalized AGC target of 500% and a maximum injection time of 7 ms.

Thermo RAW files were processed with DIA-NN against an *in-silico* library generated from the *Mus musculus* reference proteome(*88*) (UniProt ID: UP000000589). The DIA-NN output was further analyzed using FragPipe Analyst to identify differentially bound proteins(*89*). Proteomic data are deposited at MassIVE (#MSV000099437) and listed in Table S1 with file names.

### *In vivo* four-base DMS (fbDMS) probing

For endogenous mouse *Malat1* DMS probing, BV2 cells from two confluent 150mm tissue culture dishes were briefly washed with 1x PBS and then dislodged with cell scrapers in 10mL 1x PBS. After centrifugation at 1,000g for 5 min at 4°C, cell pellets were resuspended in 500 µL 1 M bicine buffer (pH 8.3) and then divided into 2 tubes with 250 µL each. The cell suspensions were either modified with 25 µL 1.7 M DMS or treated with 25 µL EtOH as a control and mixed well by pipetting. The samples were then incubated for 6 min at 37°C protected from light. For RNA extraction, 900 µL TRIzol was added into each tube. Next, 180 uL of chloroform:isoamyl alcohol (24:1) was added and incubated for 5 min and subsequently centrifuged at 3,000 g for 15 min at 4°C. The aqueous phase was then transferred to a new tube and precipitated by EtOH (at a final concentration of 70%) overnight at -20°C. The following day, RNA was pelleted at 15,000 g for 30 min at 4°C and resuspended in 1x ME buffer (10mM MOPS, 0,1mM EDTA, pH 6.5) with 1:100 RNase inhibitor. RNA was then purified with the QIAGEN RNeasy Kit (Cat. No. 74104) and eluted with 1x ME with 1:100 RNase inhibitor.

For re-probing *Malat1* IRES in the bi-cistronic reporters, pDual-GFP-(1-739) IRES-RFP or pDual-GFP-(558-695) IRES-RFP constructs were transfected into N2a cells in a 6 well plate for 48 h. Cells were washed with 1x PBS, dislodged with cell scrapers and DMS probing and RNA extraction were performed as described above.

### RT-PCR and Library Preparation

For endogenous mouse *Malat1*, a 1.1kb amplicon was designed to cover the M1 open reading frame and the upstream sequence (1-739 nt). For re-probing the *Malat1* bi-cistronic reporters, 0.8kb and 0.2kb amplicons were designed for pDual-GFP-(1-739)IRES-RFP and pDual-GFP-(558-695)IRES-RFP, respectively. Gene-specific primer sequences are listed in Table S2.

RT reactions were prepared by mixing 1 μg of total cellular RNA with 1uL of 1uM RT primer for each amplicon, annealed at 68°C for 5 min and cooled to room temperature. Next, 8uL of 2.5 x MarathonRT buffer (125mM 1M Tris-HCl pH 8.3, 500mM KCl, 12.5mM DTT, 1.25mM dNTPs, 2.5mM Mn^2+^), 4µL 100% glycerol and 0.5µL MarathonRT were added and incubated at 42°C for 3 h. The enzyme was then inactivated at 70°C for 15 min, followed by cDNA purification using AmpureXP (Cat. No. A63880) at a 1.8x bead-to-sample ratio. Purified cDNA was eluted in 10uL nuclease-free water.

*Malat1* amplicons were generated using 5 μL of cDNA and gene-specific forward and reverse primers with NEB Q5 High-Fidelity 2x Master Mix (Cat. No. M0492S) under touchdown cycling PCR conditions (68 to 58 °C annealing temperature gradient). PCR products were purified with AmpureXP (Cat. No. A63880) at a 1.8x bead-to-sample ratio and analyzed on a 1% agarose gel to verify the amplicon sizes.

For library preparation, amplicon concentrations were measured with the Qubit dsDNA HS Assay Kit (Cat No. Q32851) and amplicons diluted to 0.2 ng/μL. Libraries were generated using the Illumina NexteraXT DNA Library Preparation Kit (Cat. No. FC-131-1096) according to the manufacturer’s protocols, but at 1/5th the recommended volume. Library concentration and average fragment size were quantified with a Qubit dsDNA HS Assay Kit (ThermoFisher, Cat. No. Q32851) and a BioAnalyzer High Sensitivity DNA Analysis kit (Agilent, Cat. No. 5067-4626), respectively. Final libraries were diluted according to the manufacturer’s protocols and sequenced on an Illumina NextSeq 1000/2000 platform using a 300 cycle output kit (Cat. No. 20050264).

### fbDMS-MaP data analysis and secondary structure modelling

All sequencing data were analyzed using ShapeMapper 2.2 with DMS mode (--dms)(*36*, *90*), aligning reads to endogenous mouse *Malat1* (GenBank: 72289) or *Malat1* bi-cistronic reporter constructs (sequences listed in Table S2). Normalized reactivity values for the pDual-GFP-(558-695)IRES-RFP construct were re-scaled to match the mean of the normalized reactivities for the pDual-GFP-(1-739)IRES-RFP construct. Data that passed default quality control thresholds were used to constrain secondary structure modelling using SuperFold 1.2 with DMS mode (--DMS)(*91*). RNA secondary structures were visualized in StructureEditor(*92*).

### Data and code availability

ShapeMapper outputs and RNA secondary structure files are available at https://doi.org/10.5281/zenodo.21111515 or from the GitHub repository (https://github.com/pylelab/MALAT1-IRES-fbDMS-MaP-Structure). Raw and processed fbDMS-MaP data has been deposited on the Gene Expression Omnibus (GEO) database under the accession GSE310313.

### Plasmid constructs

For M1-GFP-Malat1 constructs, the EGFP sequence (717 bp) with removal of ATG was amplified and inserted into pSV2-Malat1 plasmid (Malat1 transcript: NR_002847) using Gibson assembly kit. The series of 100 nucleotide sequential deletions of the upstream (1-739) of M1 open reading frame were introduced in M1-GFP-Malat1 plasmids. ATG start codon of M1 ORF was mutated to TAG in M1-GFP-Malat1 plasmid for the mutation construct. The deletions and mutation were accomplished by vector PCR amplification, PNK phosphorylation and T4 ligation.

For Bicistronic reporter experiments, synthesized GFP-MCS (Bbvc1, Xba1, Ecor1 and Age1)-RFP fragment (from Twistbio) was first ligated to a linearized vector with a sv40 promoter. We termed this construct as pDual-GFP-RFP reporter. The full length of mouse MALAT1 IRES region (1-739) was amplified using forward (Ecor1 overhang) and reverse (Age1 overhang) primers. The full-length amplified IRES was ligated to Ecor1 and Age1 cleaved pDual-GFP-RFP reporter and termed as pDual-GFP-FL-IRES-RFP. A series of 100 nucleotide sequential deletions were introduced from the IRES region in pDual-GFP-IRES-RFP construct by using vector PCR amplification, PNK phosphorylation and T4 ligation method.

For Minimal IRES pDual-GFP-RFP reporter, we first amplified the minimal mouse IRES region (nucleotide 558-695) using forward (Ecor1 overhang) and reverse (Age1 overhang) primers, the amplified PCR products were then ligated to Ecor1 and Age1 linearized pDual-GFP-RFP construct using T4 ligase.

For the P1, P2, P3 deletion reporters, the whole plasmid PCR amplification was performed using forward and reverse primers to skip the deletion region, the amplified vector DNA was phosphorylated by PNK and ligated by T4 ligase.

For P1, P2, P3 insertion reporters, specific forward (Ecor1 overhang) and reverse (Age1 overhang) primers were used for amplification of different inserts. The amplified inserts were ligated to pDual-GFP-RFP vectors cleaved by Ecor1 and Age1.

### Cell transfections

For transfection of the series of M1-GFP-Malat1 plasmids, 2 ug of each construct was added to 50 ul optiMEM medium, 4 ul of lipofectamine 2000 was added to another 50 ul optiMEM. After 5 min incubation, plasmids and lipofectamine mixtures were mixed and incubated for 15 min. The ∼106 ul mixture was added into a well of 12 plates and supplemented with 400 ul fresh optiMEM. GFP fluorescent images were taken after 48 h transfection. Cells with different transfections were then harvested for WB analysis.

For transfection of a series of pDual-GFP-IRES-RFP plasmids, 2.5 ug of each construct were added to 50 ul optiMEM medium, 5 ul of lipofectamine 2000 was added to another 50 ul optiMEM. After 5 min incubation, plasmid and lipofectamine mixtures were mixed and incubated for 15 min. The mixture was then added dropwise into 12-well plates and supplemented with 400 ul fresh optiMEM. GFP and RFP fluorescent images were taken after 48 h transfection. Cells were then harvested for WB analysis using GFP, RFP and Vinculin antibodies.

For siRNA-plasmid co-transfections, 2.5 ug pDual-GFP-IRES-RFP plasmids were transfected as the method mentioned above. The siRNA transfection was performed as below: 3 ul of 50 uM siRNA was added into 50 ul optiMEM, 6 ul of lipofectamine 2000 was added into 50 ul optiMEM in another tube. After 5 min incubation, siRNA and lipofectamine mixtures were mixed and incubated for 15 min. The mixtures were then added into wells of 12-well plates together with plasmid mixtures. GFP fluorescent images were taken after 48 h transfection. Cells with different transfections were then harvested for WB analysis. For plasmid overexpression transfection, 2.5 ug pDual-GFP-IRES-RFP and 2.5 ug Rack1-Flag or 3xFlag plasmids were added into 100 ul optiMEM, 10 ul lipofectamine 2000 was added into another tube with 100 ul optiMEM. After 5 min incubation, two mixtures were mixed and further incubated for 15 min. The mixtures were added into wells of 12-well plates with 75% confluence of N2a cells. GFP and RFP fluorescent images were taken after 48 h transfection. Cells were then harvested for WB analysis using GFP, RFP and Vinculin antibodies.

### RNA extraction and RT-PCR

Cells treated with plasmids or siRNAs were collected in 500 ul TRIzol (Invitrogen, 15596018). Total RNA was isolated from aqueous phase using RNA miniprep column (Qiagen, 74106) and treated with RNase free DNase set within RDD buffer (Qiagen, 79254), followed by washing and elution according to the manufacturer’s instructions. Total RNA was reversed transcribed into cDNA with random hexamers using Superscript III Reverse Transcriptase (Thermo, 18-080-085). For reverse transcription PCR, cDNA was synthesized from 0.5 ug of total RNA. PCR was carried out using 2xgoTaq master mix with specific forward and reverse primers for gene of interest. All reactions were run for 25 cycles. PCR products were resolved by electrophoresis using 1% agarose gel. PCR primers are listed in Table S2.

### In vitro transcription and transfection

For mRNA transfection into N2a Cells, m7G or unmethylated G capped and polyadenylated mRNAs were generated by in vitro transcription. mRNAs were in vitro transcribed using the HiScript T7 kit (NEB, E2040S). A 40 ul transcription reaction contained 2 ug linearized DNA templates, 4 mM of each NTP, 3 ul of T7 RNA polymerase and 4 ul of 10 x T7 reaction buffer. For the capped RNA, GTP was substituted with 3’-O-Me-m7G(5’)ppp(5’)G RNA Cap structure analog (NEB, S1411S) in a 1:5 ratio to yield almost 75% capped RNA. For the unmethylated G capped RNA, GTP was substituted with G(5’)ppp(5’)G RNA Cap analog (NEB, S1407S) in a 1:5 ratio to yield almost 85% capped RNA. After 3 hours incubation at 37°C, 500 ul trizol was added into each reaction. The RNAs were extracted using Qiagen RNeasy mini kit (Qiagen, 74106) as mentioned in RNA extraction section.

For RNA transfection in N2a cells, 1 µg of each RNA was transfected into cells cultured in 12-well plates using Lipofectamine RNAiMAX (Ref:13778-150) per manufacturer’s instructions (Life Technologies). Media was changed 3 h after transfection and replaced with complete DMEM supplemented with 10% FBS and Pen/Strep.

For RNA aptamer (3 x S1m) affinity pulldown experiments, pcDNA3.1-3xS1m plasmids were first linearized by Asis 1 enzyme, then the linearized DNA were pelleted by ethanol and purified. RNAs were transcribed using NEB HiScribe T7 kit (NEB, E2040S) without adding cap analog. After 3-4 hours incubation at 37°C, 500 ul trizol was added in to each reaction. The RNAs were extracted using Qiagen RNA extraction column (Qiagen, 74106). RNA concentrations were measured by Nanodrop. One reaction typically yielded ∼100 ug of RNA. A small amount (∼1 ug) of each in vitro transcribed RNA was resolved in 1% agarose gel to check their quality.

### Silver staining

A series of 3xS1m-RNAs (3xS1m, 3xS1m-IRES, 3xS1m-Rev, 3xS1m-P1-del, 3xS1m-P3-del) pulled down proteins were resolved in a gradient PAGE gel and visualized by silver staining under manufacturer’s instructions (Thermo, 24612). Briefly, the PAGE gradient gel (4-12%) was washed twice with water and fixed (30% ethanol:10% acetic acid) for 30 min. The gel was washed 2 times with 10% ethanol and then 2 times with pure water. Sensitizer working solution (50μL Sensitizer with 25mL water) was added for 1 min, followed by 2 min wash with water. Stain working solution (0.5mL Enhancer with 25mL Stain) was added and incubated for 30 min. The gel was rinsed with water and then immersed in a developer working solution (0.5mL Enhancer with 25mL Developer) for 2-3 min until bands appear. The gel staining was stopped with 5% acetic acid for 10 min.

### Immunofluorescence

N2a cells were washed once with ice-cold PBSM (1xPBS, 5mM MgCl2), followed by fixation with 4% paraformaldehyde (PFA) in PBSM for 10 min at room temperature (RT). After a 5-minute wash with ice-cold PBSM, the cells were permeabilized with 0.3% Triton X-100 in PBSM for 7 min on ice. The cells were washed once with PBSM and blocked with 3% BSA (Fraction V) in PBSM for 0.5 h at room temperature. The coverslips were then incubated with primary antibody in 3% BSA in PBSM for 1 hr at RT. After 3 washes in PBST (1XPBS 0.1% tween 20), secondary antibody was diluted in 3% BSA in PBSM for 45 min at RT. Cells were washed 3 times with PBST. DAPI was added in the last wash in PBST. Cells were mounted with pro-long mounting media overnight.

### Single molecule FISH

smRNA FISH probes were house made and labeled with ATTO-647N dyes(*93*). The GFP oligo set (19 x oligos) was ordered from IDT in a wet 96-well format. Briefly, N2a cultured cells were washed once with ice-cold PBSM (1xPBS, 5mM MgCl_2_), followed by fixation with 4% paraformaldehyde in PBSM for 8 min at room temperature (RT). After a 2x wash with ice-cold PBSM, the cells were permeabilized with 70% ethanol overnight. The cells were then hydrated with 2 x SSC buffer containing 10% formamide for 1 h. RNA FISH probes were hybridized to cells at a concentration of 1 ng/ul in Hybridization buffer (Biosearch: SMF-HB1-10, 10% formamide added freshly). Coverslips were placed on a parafilm in a humidified box overnight. Cells were washed once with Wash-buffer A (Biosearch: SMF-WA1-60) at 37C for 30 min followed by washing with Wash-buffer A containing 0.5 ug/ml DAPI at 37C for another 30 min. Cells were washed with 2 x SSC buffer at RT for 5 min and mounted with 10 ul prolong anti-fade mounting media overnight at RT. The RNA smFISH probe sequences for GFP are listed in Table S2.

### Image acquisition and data analysis

The GFP and RFP fluorescent and brightfield images for bicistronic reporter assays were acquired from Nikon microscope (Nikon eclipse 2000S). For the images of the knockdown and overexpression of gene of interest for bicistronic reporter assay, GFP, RFP and DAPI fluorescence signals were acquired on a Leica sp8 light sheet confocal microscope with Z-stack. For GFP and RFP fluorescence quantification, fluorescence intensity was measured as the mean signal across the entire field of view for both GFP and RFP channels independently. For each condition, 10 randomly selected fields were acquired and the mean RFP intensity was normalized to the mean GFP intensity within the same field. All quantitative measurements were performed with imageJ imaging software(*94*). Gene ontology (GO) enrichment analysis was performed using ShinyGO (v0.85)(*95*). Volcano plots were generated using VolcaNoseR(*96*).

### Statistics

In all figures, data was presented as mean, SD or SEM as stated in the figure legend, and P < 0.05 was considered significant, ‘NS’ represents “non-significant’ for P > 0.05. *P means P<0.05, **p means p<0.01, ***p means p < 0.001, ****p means p<0.0001. P values were determined by two-tailed unpaired student’s t test unless otherwise stated, and specific p values used are indicated in the figure legends. Multiple independent experiments were performed on different ways to validate the reproductivity of experimental findings.

## Acknowledgments

We thank Dr. Fuwen Yao for assistance with heatmap analysis and Dr. Liujuan Cui for help with silver staining. We thank Rachel Green and Joel McManus for advice on IRES assays. We also thank members of the Black laboratory for their valuable advice and suggestions. Microscopy support was provided by the California NanoSystems Institute (CNSI) Imaging Core at the University of California, Los Angeles (UCLA).

## Author Contributions

W.X., A.M.P, and D.L.B. designed the research; W.X., T.H., Z.P., V.P. and J.C. performed experiments; W.X., T.H., Z.P., V.P., J.W., A.M.P, and D.L.B. analyzed the data; and W.X., T.H., and D.L.B. wrote the manuscript with input from the other authors.

## Data, Code, and Materials Availability

All data and code needed to evaluate and reproduce the results in the paper are present in the paper and/or the Supplementary Materials. Raw and processed fbDMS-MaP data have been deposited on the Gene Expression Omnibus (GEO) database under accession GSE310313 (https://www.ncbi.nlm.nih.gov/geo/query/acc.cgi?acc=GSE310313).

Proteomic data have been deposited at MassIVE under accession MSV000099437 (https://massive.ucsd.edu/ProteoSAFe/dataset.jsp?task=MSV000099437).

ShapeMapper outputs and RNA secondary structure files are available at https://doi.org/10.5281/zenodo.21111515 or from the GitHub repository (https://github.com/pylelab/MALAT1-IRES-fbDMS-MaP-Structure).

Plasmid constructs generated in this study are available from the corresponding author, Douglas L. Black, upon request.

## References

1. Y. Xiao, Y. Ren, W. Hu, A. R. Paliouras, W. Zhang, L. Zhong, K. Yang, L. Su, P. Wang, Y. Li, M. Ma, L. Shi, Long non-coding RNA-encoded micropeptides: functions, mechanisms and implications. Cell Death Discov. 10, 450 (2024).

2. J. Ruiz-Orera, X. Messeguer, J. A. Subirana, M. M. Alba, Long non-coding RNAs as a source of new peptides. eLife 3, e03523 (2014).

3. A. L. Tornesello, A. Cerasuolo, N. Starita, S. Amiranda, T. P. Cimmino, P. Bonelli, F. M. Tuccillo, F. M. Buonaguro, L. Buonaguro, M. L. Tornesello, Emerging role of endogenous peptides encoded by non-coding RNAs in cancer biology. Non-coding RNA Research 10, 231–241 (2025).

4. K. Wen, X. Chen, J. Gu, Z. Chen, Z. Wang, Beyond traditional translation: ncRNA derived peptides as modulators of tumor behaviors. J Biomed Sci 31, 63 (2024).

5. N. T. Ingolia, G. A. Brar, N. Stern-Ginossar, M. S. Harris, G. J. S. Talhouarne, S. E. Jackson, M. R. Wills, J. S. Weissman, Ribosome Profiling Reveals Pervasive Translation Outside of Annotated Protein-Coding Genes. Cell Reports 8, 1365– 1379 (2014).

6. W. Xiao, R. Halabi, C.-H. Lin, M. Nazim, K.-H. Yeom, D. L. Black, The lncRNA *Malat1* is trafficked to the cytoplasm as a localized mRNA encoding a small peptide in neurons. Genes Dev. 38, 294–307 (2024).

7. J. A. West, C. P. Davis, H. Sunwoo, M. D. Simon, R. I. Sadreyev, P. I. Wang, M. Y. Tolstorukov, R. E. Kingston, The Long Noncoding RNAs NEAT1 and MALAT1 Bind Active Chromatin Sites. Molecular Cell 55, 791–802 (2014).

8. H. Miao, F. Wu, Y. Li, C. Qin, Y. Zhao, M. Xie, H. Dai, H. Yao, H. Cai, Q. Wang, X. Song, L. Li, MALAT1 modulates alternative splicing by cooperating with the splicing factors PTBP1 and PSF. Sci. Adv. 8, eabq7289 (2022).

9. M. Huang, H. Wang, X. Hu, X. Cao, lncRNA MALAT1 binds chromatin remodeling subunit BRG1 to epigenetically promote inflammation-related hepatocellular carcinoma progression. OncoImmunology 8, e1518628 (2019).

10. G. Arun, D. Aggarwal, D. L. Spector, MALAT1 Long Non-Coding RNA: Functional Implications. ncRNA 6, 22 (2020).

11. M. Aslanzadeh, L. Stanicek, M. Tarbier, E. Mármol-Sánchez, I. Biryukova, M. R. Friedländer, Malat1 affects transcription and splicing through distinct pathways in mouse embryonic stem cells. NAR Genomics and Bioinformatics 6, lqae045 (2024).

12. S. U. Madugalle, W.-S. Liau, Q. Zhao, X. Li, H. Gong, P. R. Marshall, A. Periyakaruppiah, E. L. Zajaczkowski, L. J. Leighton, H. Ren, M. R. B. Musgrove, J. W. A. Davies, G. Kim, S. Rauch, C. He, B. C. Dickinson, B. Fulopova, L. N. Fletcher, S. R. Williams, R. C. Spitale, T. W. Bredy, Synapse-Enriched m ^6^ A-Modified Malat1 Interacts with the Novel m ^6^ A Reader, DPYSL2, and Is Required for Fear-Extinction Memory. J. Neurosci. 43, 7084–7100 (2023).

13. D. Bernard, K. V. Prasanth, V. Tripathi, S. Colasse, T. Nakamura, Z. Xuan, M. Q. Zhang, F. Sedel, L. Jourdren, F. Coulpier, A. Triller, D. L. Spector, A. Bessis, A long nuclear-retained non-coding RNA regulates synaptogenesis by modulating gene expression. EMBO J 29, 3082–3093 (2010).

14. B. Culjkovic-Kraljacic, L. Skrabanek, M. V. Revuelta, J. Gasiorek, V. H. Cowling, L. Cerchietti, K. L. B. Borden, The eukaryotic translation initiation factor eIF4E elevates steady-state m^7^ G capping of coding and noncoding transcripts. Proc. Natl. Acad. Sci. U.S.A. 117, 26773–26783 (2020).

15. J. E. Wilusz, S. M. Freier, D. L. Spector, 3′ End Processing of a Long Nuclear-Retained Noncoding RNA Yields a tRNA-like Cytoplasmic RNA. Cell 135, 919–932 (2008).

16. J. E. Wilusz, C. K. JnBaptiste, L. Y. Lu, C.-D. Kuhn, L. Joshua-Tor, P. A. Sharp, A triple helix stabilizes the 3′ ends of long noncoding RNAs that lack poly(A) tails. Genes Dev. 26, 2392–2407 (2012).

17. J. A. Brown, M. L. Valenstein, T. A. Yario, K. T. Tycowski, J. A. Steitz, Formation of triple-helical structures by the 3′-end sequences of MALAT1 and MENβ noncoding RNAs. Proc. Natl. Acad. Sci. U.S.A. 109, 19202–19207 (2012).

18. J. A. Brown, D. Bulkley, J. Wang, M. L. Valenstein, T. A. Yario, T. A. Steitz, J. A. Steitz, Structural insights into the stabilization of MALAT1 noncoding RNA by a bipartite triple helix. Nat Struct Mol Biol 21, 633–640 (2014).

19. X. Wang, C. Liu, S. Zhang, H. Yan, L. Zhang, A. Jiang, Y. Liu, Y. Feng, D. Li, Y. Guo, X. Hu, Y. Lin, P. Bu, D. Li, N6-methyladenosine modification of MALAT1 promotes metastasis via reshaping nuclear speckles. Developmental Cell 56, 702–715.e8 (2021).

20. M. Yu, Z. Cai, J. Zhang, Y. Zhang, J. Fu, X. Cui, Aberrant NSUN2-mediated m5C modification of exosomal LncRNA MALAT1 induced RANKL-mediated bone destruction in multiple myeloma. Commun Biol 7, 1249 (2024).

21. M. J. Schievelbein, C. Resende, M. M. Glennon, M. Kerosky, J. A. Brown, Global RNA modifications to the MALAT1 triple helix differentially affect thermostability and weaken binding to METTL16. Journal of Biological Chemistry 300, 105548 (2024).

22. A. Ghosh, S. P. Pandey, D. C. Joshi, P. Rana, A. H. Ansari, J. S. Sundar, P. Singh, Y. Khan, M. K. Ekka, D. Chakraborty, S. Maiti, Identification of G-quadruplex structures in MALAT1 lncRNA that interact with nucleolin and nucleophosmin. Nucleic Acids Research 51, 9415–9431 (2023).

23. T. M. Nguyen, E. B. Kabotyanski, L. C. Reineke, J. Shao, F. Xiong, J.-H. Lee, J. Dubrulle, H. Johnson, F. Stossi, P. S. Tsoi, K.-J. Choi, A. G. Ellis, N. Zhao, J. Cao, O. Adewunmi, J. C. Ferreon, A. C. M. Ferreon, J. R. Neilson, M. A. Mancini, X. Chen, J. Kim, L. Ma, W. Li, J. M. Rosen, The SINEB1 element in the long non-coding RNA Malat1 is necessary for TDP-43 proteostasis. Nucleic Acids Research 48, 2621–2642 (2020).

24. R. Miyagawa, K. Tano, R. Mizuno, Y. Nakamura, K. Ijiri, R. Rakwal, J. Shibato, Y. Masuo, A. Mayeda, T. Hirose, N. Akimitsu, Identification of *cis* - and *trans* -acting factors involved in the localization of MALAT-1 noncoding RNA to nuclear speckles. RNA 18, 738–751 (2012).

25. G. G. H. Van Den Akker, F. Zacchini, B. A. C. Housmans, L. Van Der Vloet, M. M. J. Caron, L. Montanaro, T. J. M. Welting, Current Practice in Bicistronic IRES Reporter Use: A Systematic Review. IJMS 22, 5193 (2021).

26. B. Wang, Y. Zhao, R. Li, X. Li, C. Dong, W. Mi, X. Wang, Protocol for monitoring the stability of transcription factor EB using global protein stability assay. STAR Protocols 6, 104141 (2025).

27. C. Akirtava, G. E. May, C. J. McManus, False-positive IRESes from *Hoxa9* and other genes resulting from errors in mammalian 5′ UTR annotations. Proc. Natl. Acad. Sci. U.S.A. 119, e2122170119 (2022).

28. M. Kozak, A second look at cellular mRNA sequences said to function as internal ribosome entry sites. Nucleic Acids Research 33, 6593–6602 (2005).

29. Y. Yang, Z. Wang, IRES-mediated cap-independent translation, a path leading to hidden proteome. Journal of Molecular Cell Biology 11, 911–919 (2019).

30. S. L. Miller, K. M. Green, B. Crone, J. A. Switzenberg, E. M. H. Tank, A. Krans, K. Jansen-West, C. M. Wieland, E. W. Ji, L. Petrucelli, S. J. Barmada, A. P. Boyle, P. K. Todd, Cryptic intronic transcriptional initiation generates efficient endogenous mRNA templates for C9orf72-associated RAN translation. Proc. Natl. Acad. Sci. U.S.A. 122, e2507334122 (2025).

31. B. Han, J.-T. Zhang, Regulation of Gene Expression by Internal Ribosome Entry Sites or Cryptic Promoters: the eIF4G Story. Molecular and Cellular Biology 22, 7372–7384 (2002).

32. A. G. Johnson, R. Grosely, A. N. Petrov, J. D. Puglisi, Dynamics of IRES-mediated translation. Phil. Trans. R. Soc. B 372, 20160177 (2017).

33. R. Marques, R. Lacerda, L. Romão, Internal Ribosome Entry Site (IRES)-Mediated Translation and Its Potential for Novel mRNA-Based Therapy Development. Biomedicines 10, 1865 (2022).

34. L.-T. Guo, R. L. Adams, H. Wan, N. C. Huston, O. Potapova, S. Olson, C. M. Gallardo, B. R. Graveley, B. E. Torbett, A. M. Pyle, Sequencing and Structure Probing of Long RNAs Using MarathonRT: A Next-Generation Reverse Transcriptase. Journal of Molecular Biology 432, 3338–3352 (2020).

35. R. D. C. Araujo Tavares, G. Mahadeshwar, H. Wan, A. M. Pyle, MRT-ModSeq – Rapid Detection of RNA Modifications with MarathonRT. Journal of Molecular Biology 435, 168299 (2023).

36. D. Mitchell, J. Cotter, I. Saleem, A. M. Mustoe, Mutation signature filtering enables high-fidelity RNA structure probing at all four nucleobases with DMS. Nucleic Acids Research 51, 8744–8757 (2023).

37. O. Weichenrieder, U. Kapp, S. Cusack, K. Strub, Identification of a minimal Alu RNA folding domain that specifically binds SRP9/14. RNA 3, 1262–1274 (1997).

38. O. Weichenrieder, K. Wild, K. Strub, S. Cusack, Structure and assembly of the Alu domain of the mammalian signal recognition particle. Nature 408, 167–173 (2000).

39. D. Labuda, D. Sinnett, C. Richer, J.-M. Deragon, G. Striker, Evolution of mouse B1 repeats: 7SL RNA folding pattern conserved. J Mol Evol 32, 405–414 (1991).

40. D. Labuda, E. Zitkiewicz, Evolution of secondary structure in the family of 7SL-like RNAs. J Mol Evol 39 (1994).

41. K. Leppek, G. Stoecklin, An optimized streptavidin-binding RNA aptamer for purification of ribonucleoprotein complexes identifies novel ARE-binding proteins. Nucleic Acids Research 42, e13–e13 (2014).

42. K. Leppek, K. Fujii, N. Quade, T. T. Susanto, D. Boehringer, T. Lenarčič, S. Xue, N. R. Genuth, N. Ban, M. Barna, Gene- and Species-Specific Hox mRNA Translation by Ribosome Expansion Segments. Molecular Cell 80, 980–995.e13 (2020).

43. Y. Levin, S. Bahn, “Quantification of Proteins by Label-Free LC-MS/MS” in LC-MS/MS in Proteomics, P. R. Cutillas, J. F. Timms, Eds. (Humana Press, Totowa, NJ, 2010; http://link.springer.com/10.1007/978-1-60761-780-8_13)vol. 658 of *Methods in Molecular Biology*, pp. 217–231.

44. S. Gunaseelan, K. Z. Wong, N. Min, J. Sun, N. K. B. M. Ismail, Y. J. Tan, R. C. H. Lee, J. J. H. Chu, Prunin suppresses viral IRES activity and is a potential candidate for treating enterovirus A71 infection. Sci. Transl. Med. 11, eaar5759 (2019).

45. J. Zhu, A. Korostelev, D. A. Costantino, J. P. Donohue, H. F. Noller, J. S. Kieft, Crystal structures of complexes containing domains from two viral internal ribosome entry site (IRES) RNAs bound to the 70S ribosome. Proc. Natl. Acad. Sci. U.S.A. 108, 1839–1844 (2011).

46. C. S. Fraser, J. A. Doudna, Structural and mechanistic insights into hepatitis C viral translation initiation. Nat Rev Microbiol 5, 29–38 (2007).

47. J. S. Kieft, K. Zhou, R. Jubin, J. A. Doudna, Mechanism of ribosome recruitment by hepatitis C IRES RNA. RNA 7, 194–206 (2001).

48. C. M. T. Spahn, J. S. Kieft, R. A. Grassucci, P. A. Penczek, K. Zhou, J. A. Doudna, J. Frank, Hepatitis C Virus IRES RNA-Induced Changes in the Conformation of the 40 *S* Ribosomal Subunit. Science 291, 1959–1962 (2001).

49. K. Majzoub, M. L. Hafirassou, C. Meignin, A. Goto, S. Marzi, A. Fedorova, Y. Verdier, J. Vinh, J. A. Hoffmann, F. Martin, T. F. Baumert, C. Schuster, J.-L. Imler, RACK1 Controls IRES-Mediated Translation of Viruses. Cell 159, 1086–1095 (2014).

50. J. Nilsson, J. Sengupta, J. Frank, P. Nissen, Regulation of eukaryotic translation by the RACK1 protein: a platform for signalling molecules on the ribosome. EMBO Reports 5, 1137–1141 (2004).

51. H. D. Kim, E. Kong, Y. Kim, J.-S. Chang, J. Kim, RACK1 depletion in the ribosome induces selective translation for non-canonical autophagy. Cell Death Dis 8, e2800–e2800 (2017).

52. S. Gallo, S. Ricciardi, N. Manfrini, E. Pesce, S. Oliveto, P. Calamita, M. Mancino, E. Maffioli, M. Moro, M. Crosti, V. Berno, M. Bombaci, G. Tedeschi, S. Biffo, RACK1 Specifically Regulates Translation through Its Binding to Ribosomes. Molecular and Cellular Biology 38, e00230–18 (2018).

53. Y. Jung, J.-Y. Seo, H. G. Ryu, D.-Y. Kim, K.-H. Lee, K.-T. Kim, BDNF-induced local translation of *GluA1* is regulated by HNRNP A2/B1. Sci. Adv. 6, eabd2163 (2020).

54. L. Li, X. Li, H. Zhong, M. Li, B. Wan, W. He, Y. Zhang, Y. Du, D. Chen, W. Zhang, P. Ji, D. Jiang, S. Han, VP3 protein of Senecavirus A promotes viral IRES-driven translation and attenuates innate immunity by specifically relocalizing hnRNPA2B1. J Virol 98, e01227–24 (2024).

55. C.-Y. Hung, Y.-C. Wang, J.-Y. Chuang, M.-J. Young, H. Liaw, W.-C. Chang, J.-J. Hung, Nm23-H1-stabilized hnRNPA2/B1 promotes internal ribosomal entry site (IRES)-mediated translation of Sp1 in the lung cancer progression. Sci Rep 7, 9166 (2017).

56. Y. Liu, S. Shi, The roles of HNRNP A2 / B1 in RNA biology and disease. WIREs RNA 12, e1612 (2021).

57. E. Martinez-Salas, R. Francisco-Velilla, J. Fernandez-Chamorro, A. M. Embarek, Insights into Structural and Mechanistic Features of Viral IRES Elements. Front. Microbiol. 8, 2629 (2018).

58. S. Weingarten-Gabbay, S. Elias-Kirma, R. Nir, A. A. Gritsenko, N. Stern-Ginossar, Z. Yakhini, A. Weinberger, E. Segal, Systematic discovery of cap-independent translation sequences in human and viral genomes. Science 351, aad4939 (2016).

59. S. W. Abdullah, J. Wu, X. Wang, H. Guo, S. Sun, Advances and Breakthroughs in IRES-Directed Translation and Replication of Picornaviruses. mBio 14, e00358–23 (2023).

60. N. Chamond, J. Deforges, N. Ulryck, B. Sargueil, 40S recruitment in the absence of eIF4G/4A by EMCV IRES refines the model for translation initiation on the archetype of Type II IRESs. Nucleic Acids Research 42, 10373–10384 (2014).

61. I. B. Lomakin, C. U. T. Hellen, T. V. Pestova, Physical Association of Eukaryotic Initiation Factor 4G (eIF4G) with eIF4A Strongly Enhances Binding of eIF4G to the Internal Ribosomal Entry Site of Encephalomyocarditis Virus and Is Required for Internal Initiation of Translation. Molecular and Cellular Biology 20, 6019–6029 (2000).

62. S. L. De Quinto, E. Lafuente, E. Martínez-Salas, IRES interaction with translation initiation factors: Functional characterization of novel RNA contacts with eIF3, eIF4B, and eIF4GII. RNA 7, 1213–1226 (2001).

63. J. Mailliot, F. Martin, Viral internal ribosomal entry sites: four classes for one goal. WIREs RNA 9, e1458 (2018).

64. V. Ahl, H. Keller, S. Schmidt, O. Weichenrieder, Retrotransposition and Crystal Structure of an Alu RNP in the Ribosome-Stalling Conformation. Molecular Cell 60, 715–727 (2015).

65. N. Zhu, M. Ahmed, Y. Li, J. C. Liao, P. K. Wong, Long noncoding RNA MALAT1 is dynamically regulated in leader cells during collective cancer invasion. Proc. Natl. Acad. Sci. U.S.A. 120, e2305410120 (2023).

66. W. Barczak, S. M. Carr, G. Liu, S. Munro, A. Nicastri, L. N. Lee, C. Hutchings, N. Ternette, P. Klenerman, A. Kanapin, A. Samsonova, N. B. La Thangue, Long non-coding RNA-derived peptides are immunogenic and drive a potent anti-tumour response. Nat Commun 14, 1078 (2023).

67. B. Bujisic, H.-G. Lee, L. Xu, U. Weissbein, C. Rivera, I. Topisirovic, J. T. Lee, 7SL RNA and signal recognition particle orchestrate a global cellular response to acute thermal stress. Nat Commun 16, 1630 (2025).

68. E. Ullu, C. Tschudi, Alu sequences are processed 7SL RNA genes. Nature 312, 171–172 (1984).

69. A. K. K. Lakkaraju, C. Mary, A. Scherrer, A. E. Johnson, K. Strub, SRP Keeps Polypeptides Translocation-Competent by Slowing Translation to Match Limiting ER-Targeting Sites. Cell 133, 440–451 (2008).

70. Y. Lustig, H. Goldshmidt, S. Uliel, S. Michaeli, The *Trypanosoma brucei* signal recognition particle lacks the Alu-domain-binding proteins: purification and functional analysis of its binding proteins by RNAi. Journal of Cell Science 118, 4551–4562 (2005).

71. M. Halic, T. Becker, M. R. Pool, C. M. T. Spahn, R. A. Grassucci, J. Frank, R. Beckmann, Structure of the signal recognition particle interacting with the elongation-arrested ribosome. Nature 427, 808–814 (2004).

72. B. Beckert, A. Kedrov, D. Sohmen, G. Kempf, K. Wild, I. Sinning, H. Stahlberg, D. N. Wilson, R. Beckmann, Translational arrest by a prokaryotic signal recognition particle is mediated by RNA interactions. Nat Struct Mol Biol 22, 767–773 (2015).

73. A. Berger, E. Ivanova, C. Gareau, A. Scherrer, R. Mazroui, K. Strub, Direct binding of the Alu binding protein dimer SRP9/14 to 40S ribosomal subunits promotes stress granule formation and is regulated by Alu RNA. Nucleic Acids Research 42, 11203–11217 (2014).

74. D. E. A. Birse, U. Kapp, K. Strub, S. Cusack, A. Åberg, The crystal structure of the signal recognition particle Alu RNA binding heterodimer, SRP9/14. EMBO J 16, 3757–3766 (1997).

75. J. Hasler, Alu RNP and Alu RNA regulate translation initiation in vitro. Nucleic Acids Research 34, 2374–2385 (2006).

76. J. B. Moldovan, H. C. Kopera, Y. Liu, M. Garcia-Canadas, P. Catalina, P. E. Leone, L. Sanchez, J. O. Kitzman, J. M. Kidd, J. L. Garcia-Perez, J. V. Moran, Variable patterns of retrotransposition in different HeLa strains provide mechanistic insights into SINE RNA mobilization processes. Nucleic Acids Research 52, 7761–7779 (2024).

77. E. Sundaramoorthy, M. Leonard, R. Mak, J. Liao, A. Fulzele, E. J. Bennett, ZNF598 and RACK1 Regulate Mammalian Ribosome-Associated Quality Control Function by Mediating Regulatory 40S Ribosomal Ubiquitylation. Molecular Cell 65, 751–760.e4 (2017).

78. I. A. Muslimov, A. Tuzhilin, T. H. Tang, R. K. S. Wong, R. Bianchi, H. Tiedge, Interactions of noncanonical motifs with hnRNP A2 promote activity-dependent RNA transport in neurons. Journal of Cell Biology 205, 493–510 (2014).

79. J. Shan, T. P. Munro, E. Barbarese, J. H. Carson, R. Smith, A Molecular Mechanism for mRNA Trafficking in Neuronal Dendrites. J. Neurosci. 23, 8859– 8866 (2003).

80. J. Lo, K. F. Vaeth, G. Bhardwaj, N. Mukherjee, H. A. Russ, J. K. Moore, J. M. Taliaferro, The RNA binding protein HNRNPA2B1 regulates RNA abundance and motor protein activity in neurites. [Preprint] (2024). 10.1101/2024.08.26.609768.

81. H.-J. Kim, H.-R. Lee, J.-Y. Seo, H. G. Ryu, K.-H. Lee, D.-Y. Kim, K.-T. Kim, Heterogeneous nuclear ribonucleoprotein A1 regulates rhythmic synthesis of mouse Nfil3 protein via IRES-mediated translation. Sci Rep 7, 42882 (2017).

82. M. Charpentier, E. Dupré, A. Fortun, F. Briand, M. Maillasson, E. Com, C. Pineau, N. Labarrière, C. Rabu, F. Lang, hnRNP-A1 binds to the IRES of MELOE-1 antigen to promote MELOE-1 translation in stressed melanoma cells. Molecular Oncology 16, 594–606 (2022).

83. J. Feng, J. Zhou, Y. Lin, W. Huang, hnRNP A1 in RNA metabolism regulation and as a potential therapeutic target. Front. Pharmacol. 13, 986409 (2022).

84. S. Braunstein, K. Karpisheva, C. Pola, J. Goldberg, T. Hochman, H. Yee, J. Cangiarella, R. Arju, S. C. Formenti, R. J. Schneider, A Hypoxia-Controlled Cap-Dependent to Cap-Independent Translation Switch in Breast Cancer. Molecular Cell 28, 501–512 (2007).

85. K. M. Bedard, S. Daijogo, B. L. Semler, A nucleo-cytoplasmic SR protein functions in viral IRES-mediated translation initiation. EMBO J 26, 459–467 (2007).

86. E. Blasi, R. Barluzzi, V. Bocchini, R. Mazzolla, F. Bistoni, Immortalization of murine microglial cells by a v-raf / v-myc carrying retrovirus. Journal of Neuroimmunology 27, 229–237 (1990).

87. T. S. Batth, MaximA. X. Tollenaere, P. Rüther, A. Gonzalez-Franquesa, B. S. Prabhakar, S. Bekker-Jensen, A. S. Deshmukh, J. V. Olsen, Protein Aggregation Capture on Microparticles Enables Multipurpose Proteomics Sample Preparation*. Molecular & Cellular Proteomics 18, 1027a–11035 (2019).

88. V. Demichev, C. B. Messner, S. I. Vernardis, K. S. Lilley, M. Ralser, DIA-NN: neural networks and interference correction enable deep proteome coverage in high throughput. Nat Methods 17, 41–44 (2020).

89. Y. Hsiao, H. Zhang, G. X. Li, Y. Deng, F. Yu, H. V. Kahrood, J. R. Steele, R. B. Schittenhelm, A. I. Nesvizhskii, Analysis and visualization of quantitative proteomics data using FragPipe-Analyst. [Preprint] (2024). 10.1101/2024.03.05.583643.

90. S. Busan, K. M. Weeks, Accurate detection of chemical modifications in RNA by mutational profiling (MaP) with ShapeMapper 2. RNA 24, 143–148 (2018).

91. M. J. Smola, G. M. Rice, S. Busan, N. A. Siegfried, K. M. Weeks, Selective 2′-hydroxyl acylation analyzed by primer extension and mutational profiling (SHAPE-MaP) for direct, versatile and accurate RNA structure analysis. Nat Protoc 10, 1643–1669 (2015).

92. J. S. Reuter, D. H. Mathews, RNAstructure: software for RNA secondary structure prediction and analysis. BMC Bioinformatics 11, 129 (2010).

93. W. Xiao, K.-H. Yeom, C.-H. Lin, D. L. Black, Improved enzymatic labeling of fluorescent in situ hybridization probes applied to the visualization of retained introns in cells. RNA 29, 1274–1287 (2023).

94. C. A. Schneider, W. S. Rasband, K. W. Eliceiri, NIH Image to ImageJ: 25 years of image analysis. Nat Methods 9, 671–675 (2012).

95. S. X. Ge, D. Jung, R. Yao, ShinyGO: a graphical gene-set enrichment tool for animals and plants. Bioinformatics 36, 2628–2629 (2020).

96. J. Goedhart, M. S. Luijsterburg, VolcaNoseR is a web app for creating, exploring, labeling and sharing volcano plots. Sci Rep 10, 20560 (2020).

